# LncRNA-mediated organization of oncogenic chromatin state is a targetable vulnerability in leukemia

**DOI:** 10.64898/2026.08.10.744058

**Authors:** Zhen Jin, Brenda Ma, Karen Y.T. Chan, Diana Lin, Debajeet Ghosh, Amber Louwagie, Luke Saville, Joanne Liani Chandra, Jennifer Grants, Yilin Liu, Zongmin Liu, Leo Escano, Se-Wing Grace Cheng, Sandra E. Spencer, Priscila E. F. Stricker, Glenn Edin, Fiona Wong, Kevin Dong, Quang Anh Hoang, Quang Thang T. Bui, Alexandra Schurer, Kim Do, Tim Chou, Oakes Christopher, Genc Basha, Martin Sauvageau, Samer Hussein, Gregg B. Morin, Fabiana Perna, Florian Kuchenbauer, Michael G. Kharas, Aly Karsan, Pieter R. Cullis, Ly P. Vu

## Abstract

Long non-coding RNAs (lncRNAs) are increasingly recognized as critical regulators of gene expression underlying various cellular functions, however, the functional and mechanistic contributions of most lncRNAs to tumorigenesis remain poorly defined, and targeting of lncRNAs is challenging with conventional therapeutic approaches. Here, we uncover human *PAN3-AS1* and its murine ortholog *Lnc35682*, previously uncharacterized lncRNAs embedded within a conserved syntenic genomic locus, as highly expressed in acute myeloid leukemia (AML). Using genetic mouse models, human cell lines and primary patient samples, we show that *PAN3-AS1* is essential for leukemia maintenance but dispensable for normal hematopoiesis. Mechanistically, we find that elevated *PAN3-AS1* influences chromatin accessibility, thus promoting leukemia gene expression programs. This is mediated, at least in part, by *PAN3-*AS1’s association with the nuclear lamina through a defined functional region that is required for its leukemogenic function. We further characterize a feed-forward regulatory circuit between *PAN3-AS1* and its neighboring gene *FLT3* that directly links the aberrant lncRNA functions to the FLT3-mutant AML subtype. To therapeutically exploit the regulatory node, we engineer a myeloid leukemia-preferentially targeted lipid nanoparticle (LNP) formulation and demonstrate effective delivery of siRNAs against endogenous targets into leukemia cells in experimental animals. LNP-siPAN3-AS1 alone or in combination with a clinically used FLT3 inhibitor, Gilteritinib, reduces leukemia burden and significantly delay leukemogenesis *in vivo*. Overall, our study uncovers a therapeutic vulnerable lncRNA-centric circuitry and provides compelling preclinical evidence for the development and application of a novel RNA targeting-LNP based therapy for treatment of myeloid leukemia.

**Graphical abstract:** 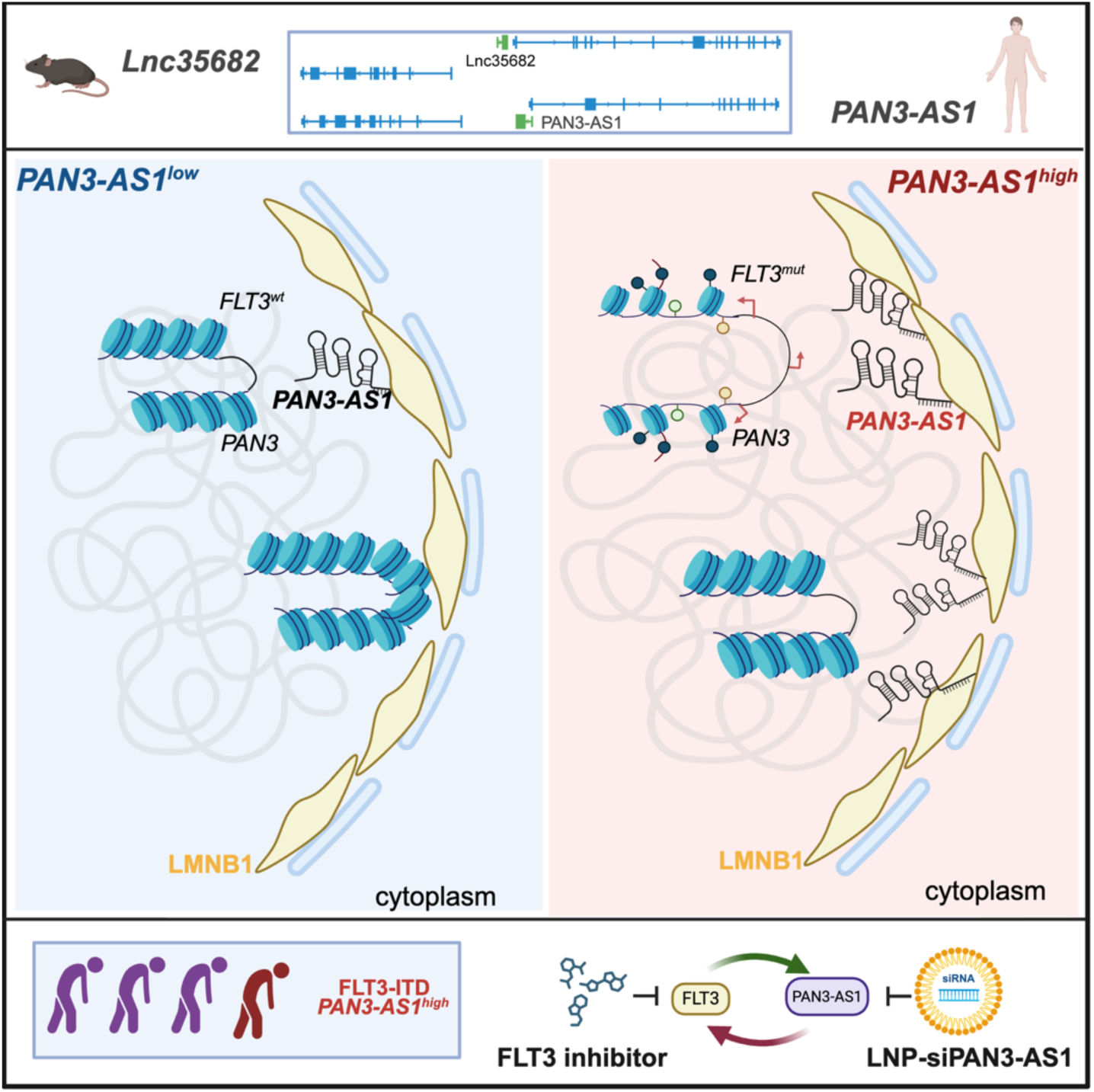

## Introduction

Dysregulation of gene expression leads to disruption of normal development and tumorigenesis. It has been widely established that only 2% of the human genome encodes for proteins and a large portion of the genome is transcribed but not translated (Djebali et al., 2012). The discovery of long non-coding RNAs (lncRNAs) among transcriptional products from non-coding regions of the genomes have expanded the catalog of regulatory molecules that can influence gene expression. Long non-coding RNAs (lncRNAs), broadly defined as RNA transcripts of greater than 200 nucleotides without protein coding potential, constitute of heterogenous groups of diverse non-coding transcripts including sense, antisense, intronic or intergenic to protein-coding loci (Morris and Mattick, 2014). LncRNAs are versatile molecules, capable of binding to DNA, RNA and proteins (Rinn and Chang, 2012), allowing lncRNAs to regulate gene expression at multiple levels (Statello et al., 2021). Given their intricate involvement in gene expression control, lncRNAs have been shown to modulate diverse cellular processes such as cell pluripotency, differentiation and apoptosis, hence contributing directly to many aspects of cancer hallmarks (Schmitt and Chang, 2016). LncRNA expression profiles can be used to classify clinically relevant cancer subtypes and predict disease prognosis (Du et al., 2013). Mutations and aberrant expression of lncRNAs have been identified in several cancer types including leukemia, lung, breast, prostate and gastric cancer etc (Carlevaro-Fita et al., 2020; Liu et al., 2021). However, while the annotation and general importance of lncRNAs in cancer is well accepted, only handful have been functionally characterized and mechanistically studied (Statello et al., 2021). The biological relevance and precise function of the vast majority of lncRNAs remains to be elucidated.

Despite lncRNAs’ intriguing biology, lncRNAs are frequently dismissed as therapeutic targets and efforts to leverage unique features of lncRNAs for therapeutic interventions is limited (Winkle et al., 2021). In addition to a lack of functional studies, there are several substantial barriers in translating the therapeutic potential of lncRNAs to clinical applications. Due to their complex structures and highly context-dependent functions, lncRNAs are generally considered “undruggable” by traditional drug targeting approaches. While gene silencing using siRNAs or anti-sense oligonucleotides (ASOs) can be used to perturb lncRNA expression in the laboratory, the delivery of these molecules to specific human organs poses a major hurdle (Esmaeilpour et al., 2025). Besides, as lncRNAs are often poorly conserved across species, *in vivo* pre-clinical studies of lncRNAs are not compatible with common murine disease models and instead, require the establishment of appropriate human patient-derived xenograft (PDX) models. Therefore, the key to advancing the field lies in identification of novel, therapeutically actionable lncRNA targets, successful development of effective strategies to modulate their activity *in vivo*, and establishment of delivery platforms capable of supporting RNA-based therapies.

Acute myeloid leukemia (AML) is a rapidly progressing hematological malignancy characterized by uncontrolled expansion of differentiation-arrest leukemia blasts. AML is aggressive and often fatal with a 5-year overall survival of ∼30% (Shimony et al., 2025). With the use of genomic tools in cancer diagnosis and classification, it is now well established that AML represents a group of genetically heterogenous diseases harboring recurrent chromosomal translocations e.g. t(9;11)/*MLL-AF9* rearrangement; inv(3)/*MECOM* (*EVI1*) and mutations such as *TP53, DNMT3A*, *TET2*, IDH1/2, NPM1 and *FLT3* (Do€hner et al., 2022; Papaemmanuil et al., 2016). In recent years, evidence of a role for lncRNAs in hematological malignancies has emerged (Connerty and Lock, 2023; Gourvest et al., 2019). Large scale genomic analysis showed distinct lncRNAs expression signatures associated with AML subtypes (Garzon et al., 2014), treatment responses (De Clara et al., 2017), and patient prognosis (Dohner et al., 2017). A number of lncRNAs have been characterized as oncogenes e.g., lncRNA-H19 (Zhang et al., 2018), HOTAIR (Zhang et al., 2016) or tumor suppressors e.g., NEAT1 (Zeng et al., 2014), and MEG3 (Lyu et al., 2017). However, these studies only captured a small fraction of this regulatory class and to date, no lncRNA-targeting therapy has yet been clinically investigated in myeloid leukemia.

In this study, we report the identification of human lncRNA *PAN3-AS1* and its murine ortholog *Lnc35682* as a leukemia-promoting lncRNA and establish an effective RNA interference (siRNA)-lipid nanoparticle (LNP)-based therapeutic approach targeting *PAN3-AS1* to suppress leukemia progression in primary human PDX models. We demonstrated that depletion of *PAN3-AS1* and *Lnc35682* suppresses leukemia cell proliferation and leukemogenesis *in vivo* while having minimal effects on function of normal human and mouse hematopoietic stem/progenitor cells (HSPCs). We found that *PAN3-AS1* expression is elevated in leukemia cells and its high expression can drive aggressive disease progression. Mechanistically, we discovered that *PAN3-AS1* facilitates chromatin accessibility at neighboring genes, including the proto-oncogene FLT3, as well as at other genomic regions associated with leukemia-promoting gene expression programs. We uncovered a reciprocal regulation loop between *PAN3-AS1* and *FLT3*, which provides an exquisite targetable vulnerability for FLT3 mutant AMLs. The leukemia-specific LNP formulation enabled delivery of siRNA targeting *PAN3-AS1* in animal models and demonstrated significant disease control. Together, our work highlights lncRNAs as critical regulators of leukemogenesis and a promising therapeutic class. We provide compelling preclinical data supporting the utility of novel LNP delivery platform and RNA-based therapeutics in treatment of leukemia. Our study represents an experimental framework to uncover and characterize novel non-coding RNA regulators of malignancies and to develop effective strategy to translate these fundamental discoveries to potential therapy.

## Results

### Identification of human lncRNA *PAN3-AS1*, a leukemia-essential lncRNA embedded in a conserved genomic region

To select candidate lncRNAs for experimental investigation in AML, we developed a criteria-based framework to navigate the thousands of lncRNAs identified in the human genome that remain functionally uncharacterized. We prioritized a lncRNA candidate based on: (i) its relative abundance; (ii) an expression profile exhibiting a selectively high level in AML versus healthy counterparts and in leukemia stem cells (LSCs) versus blasts; (iii) AML specificity compared with other tissues and cancer types; and (iv) conservation in its genomic location between murine and human homologs to account for lncRNAs’ limited sequence conservation among species (Quinn et al., 2016). To obtain a paired comparison between normal HSPCs vs. LSCs, we utilized the murine MLL-AF9 leukemia model where leukemia can be generated from direct transformation of HSPCs by oncogene MLL-AF9 (Cozzio et al., 2003). In this model, LSCs have also been well phenotypically and functionally defined to be enriched in the c-kit high population of leukemia blasts (Somervaille and Cleary, 2006). Through the survey, we identified the human lncRNA *PAN3-AS1*, and the murine unannotated lncRNA at the genomic locus D5Ert605e, which we named *Lnc35682*. *PAN3-AS1* and *Lnc35682* reside within a conserved syntenic genomic block containing *FLT3* and *PAN3* on human chromosome 13 and murine chromosome 5 (**Figure 1A**). *PAN3-AS1* is highly upregulated in AML cells compared to normal hematopoietic cells (**Figure 1B**). *PAN3-AS1* is most highly elevated in AML in comparison to other cancer types, indicating high specificity for its function in AML (**Figure S1A**). On the other hand, *Lnc35682* is highly upregulated in MF9-LSCs compared to normal stem and progenitor cells (HSC, MPP, CMP, GMP, MEP), or differentiated cells (Myeloid, B cells, T cells) (**Figure 1C**). Moreover, high *PAN3-AS1* correlates with poor outcomes of AML patients, suggesting an important role in AML pathogenesis (**Figure 1D**). Given these features, we chose *PAN3-AS1* and its syntenic murine ortholog *Lnc35682* for further investigations.

**Figure 1.**
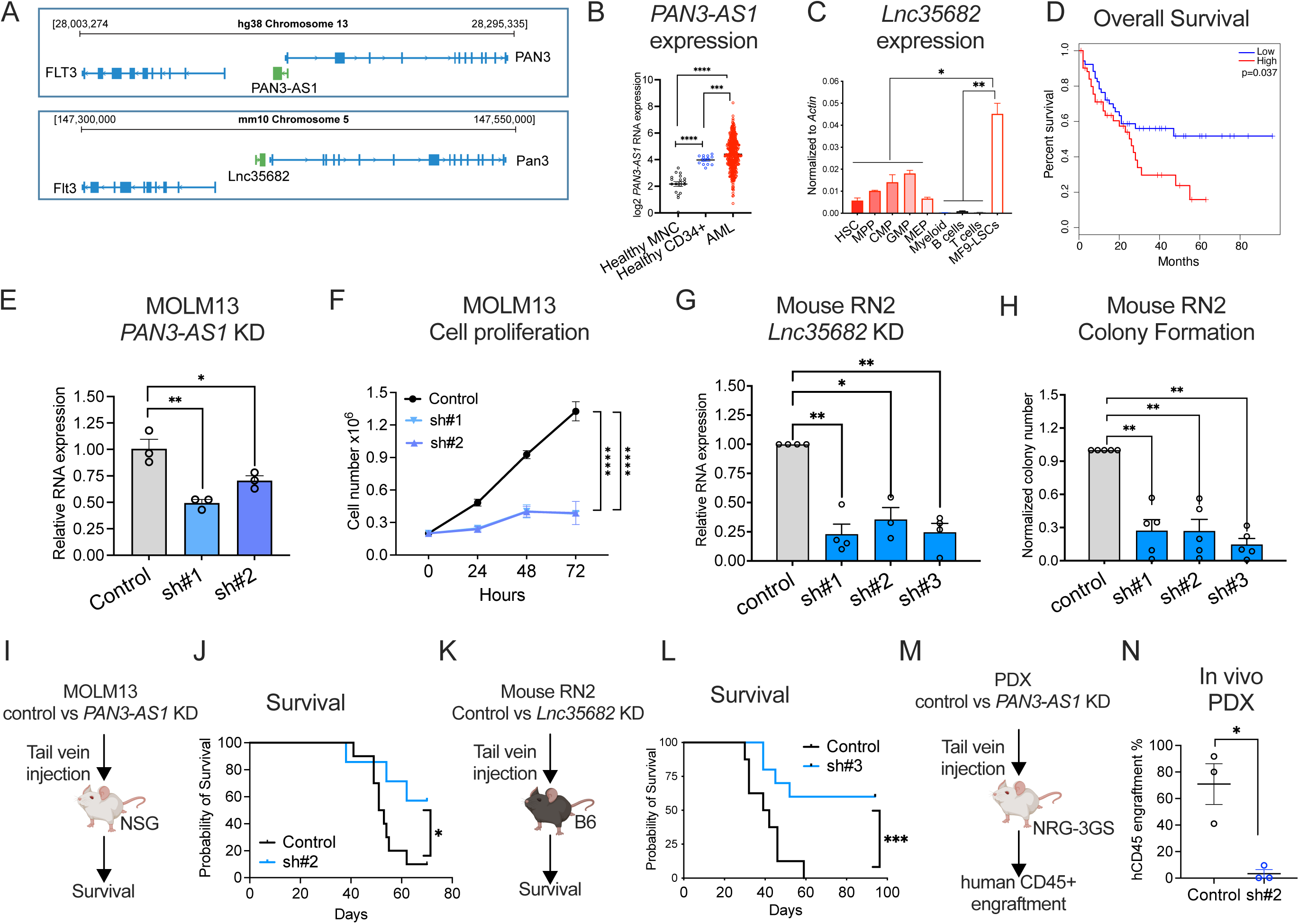
The syntenically conserved lncRNA *PAN3-AS1* is essential for myeloid leukemia. (A) Scheme of the syntenically conserved gene blocks containing lncRNA *PAN3-AS1* on human chromosomal 13 and *Lnc35682* on the murine chromosome 5. (B) Relative *PAN3-AS1* gene expression levels in healthy mononuclear cells (MNC); CD34+ enriched HSPC cells and AML cells in the BeatAML clinical cohort. MNC (N=19), CD34+ (n=13), AML (n=451). *** p<0.001, **** p<0.0001, two-tailed Student’s t-test. (C) Quantitative RT-qPCR analysis of *Lnc35682*’s expression levels in murine hematopoietic compartments. HSC, hematopoietic stem cell; MPP, multipotent progenitor; CMP, common myeloid progenitor; GMP, granulocyte monocyte progenitor; MEP, megakaryocyte erythrocyte progenitor; myeloid; B cells and T cells and ckit+ enriched LSC-dsRed leukemia cells. n=4/cell type. Data shown, mean ± s.e.m., p-Value: *<0.05, **<0.01, two-tailed Student’s t-test. (D) High-level *PAN3-AS1* expression correlated with poor prognosis of AML. Kaplan–Meier curves showing outcomes of AML patients in TCGA-AML high *PAN3-AS1* n=52 vs. low *PAN3-AS1* n=52. (E) Quantitative RT-qPCR analysis of *PAN3-AS1* expression levels in MOLM13 expressing either scrambled shRNA (control) or shRNAs targeting *PAN3-AS1* (sh#1 and sh#2). n=3. Data shown, mean ± s.e.m., p-Value: *<0.05, **<0.01, two-tailed Student’s t-test. (F) Cell proliferation of MOLM13 described in (E). n=3. Data shown, p-Value ****<0.0001, Two-way ANOVA. (G) Quantitative RT-qPCR analysis of *Lnc35682* expression in murine RN2 leukemia following retroviruses transduction of vectors carrying either scrambled shRNA (control) or shRNAs to knockdown *Lnc35682* (KD, sh#1, #2, #3). (H) Colony formation of the murine RN2 leukemia cells described in (G). (I) Experimental scheme for assessing leukemia progression *in vivo* after transplantation of MOLM13 cells transduced with control vs. sh#2 targeting *PAN3-AS1*. (J) Kaplan–Meier survival analysis of mice described in (I). Control, n = 10; knockdown, n = 7. p-Value: *<0.05, log-rank test. (K) Experimental scheme for assessing leukemia progression *in vivo* after transplantation of murine RN2 cells transduced with control vs. sh#3 targeting *Lnc35682*. (L) Kaplan–Meier survival analysis of mice described in (K). Control, n = 8; knockdown, n = 10. p-Value: ***<0.001, log-rank test. (M) Experimental scheme for assessing leukemia progression *in vivo* after transplantation of patient derived leukemia cells (PDX) transduced with control vs. sh#2 targeting *PAN3-AS1*. (N) Percentage of human leukemia CD45+ cells in recipient animals 12 weeks post transplantation. Data shown, mean ± s.e.m., p-Value: *<0.05, two-tailed Student’s t-test.

We first screened the expression levels of *PAN3-AS1* across different AML cell lines to confirm its upregulation and select cell lines with high *PAN3-AS1* expression levels for functional assessments (**Figure S1B**). We then knockdown (KD) *PAN3-AS1* using lentiviral-shRNAs (scrambled-control vs. sh#1 and sh#2 targeting *PAN3-AS1*) in MOLM13, MV4-11 and OCI-AML3 leukemia cells. We observed significant reduction in cell proliferation in all three cell lines (**Figures 1E-F and S1C-D**). To complement the shRNA-mediated KD approach, we performed sgRNA-mediated depletion of *PAN3-AS1* in MV4-11 line constitutively expressing Cas13Rx (Wessels et al., 2020) (**Figure S1E-F**). Despite a less efficient downregulation of *PAN3-AS1* expression (**Figure S1G**), we still observed an inhibition in cell growth (**Figures S1H**). In parallel, depletion of *Lnc35682* using shRNA knockdown (scrambled-control vs. sh#1, sh#2 and sh#3 targeting *Lnc35682*) markedly reduced colony forming ability of murine RN2 leukemia cells (Zuber et al., 2011) (**Figures 1G-H**). *In vivo* xenograft model showed mice receiving *PAN3-AS1* KD MOLM13 cells had lower disease burden and improved survival outcome (**Figures 1I-J and S1I**). Consistently, *Lnc35682* depletion also resulted in delayed leukemia development (**Figure 1K-L**). In patient derived leukemia cells, *PAN3-AS1* depletion inhibited leukemia cell colony formation *in vitro* (**Figure S1J**) and reduced leukemia engraftment burden in immune-deficient NRG-3GS mice (**Figure 1M-N**). These results demonstrate that *PAN3-AS1* is essential for AML cell survival and maintenance of leukemogenesis *in vivo*.

### Human *PAN3-AS1* and mouse *Lnc35682* exhibits conserved leukemia promoting activity

To further investigate *PAN3-AS1* and *Lnc35682* cellular function, we performed fluorescence *in situ* hybridization (FISH) using RNA scope to characterize their subcellular localization in AML cells. To enhance detectability and validate RNA probe specificity, we synthesized and exogenously overexpressed (OV) cDNAs encoding *PAN3-AS1* and *Lnc35682* in MOLM13 and RN2 cells. We observed that in both parental and OV cell lines, *PAN3-AS1* and *Lnc35682* exhibited a punctate perinuclear localization pattern (**Figures 2A-B and S2A**). A similar pattern for endogenous *PAN3-AS1* is noted in patient AML cells (**Figure 2C**). Next, we asked whether elevated *PAN3-AS1* and *Lnc35682* expression, as observed in AML patients, can promote AML aggressiveness. We found that overexpression of *PAN3-AS1* strongly promoted growth of MOLM13 cells (**Figure 2D and S2B**). Exogenous expression of *PAN3-AS1* also efficiently rescued the growth defects cause by *PAN3-AS1* shRNA mediated knockdown, confirming the specificity of shRNAs in targeting *PAN3-AS1* and the observed biological phenotypes (**Figures S2D and S2E**). Furthermore, mice transplanted with *PAN3-AS1*–overexpressing cells showed higher disease burden and accelerated disease progression in the *in vivo* MOLM13 xenograft (**Figures 2E, 2F and S2F**). Moreover, *Lnc35682* overexpression increased proliferation and colony forming ability of murine RN2 leukemia cells (**Figures 2G-H and S2C**). To test whether *Lnc35682* presence can mimic *PAN3-AS1*, we expressed *Lnc35682* in human MOLM13 cells. Interestingly, even though forced expression of the murine *Lnc35682* did not affect the endogenous level of *PAN3-AS1*, it enhanced proliferation of MOLM13 cells (**Figures S2G-H**). More importantly, we observed that *Lnc35682* rescued the growth defects caused by *PAN3-AS1* depletion, suggesting that there is, at least, a partial functional compensation between *PAN3-AS1* and *Lnc35682* (**Figure 2I**). Together, these findings indicate a conserved oncogenic function of the human lncRNA *PAN3-AS1* and mouse *Lnc25682* in promoting AML survival and progression when upregulated.

**Figure 2.**
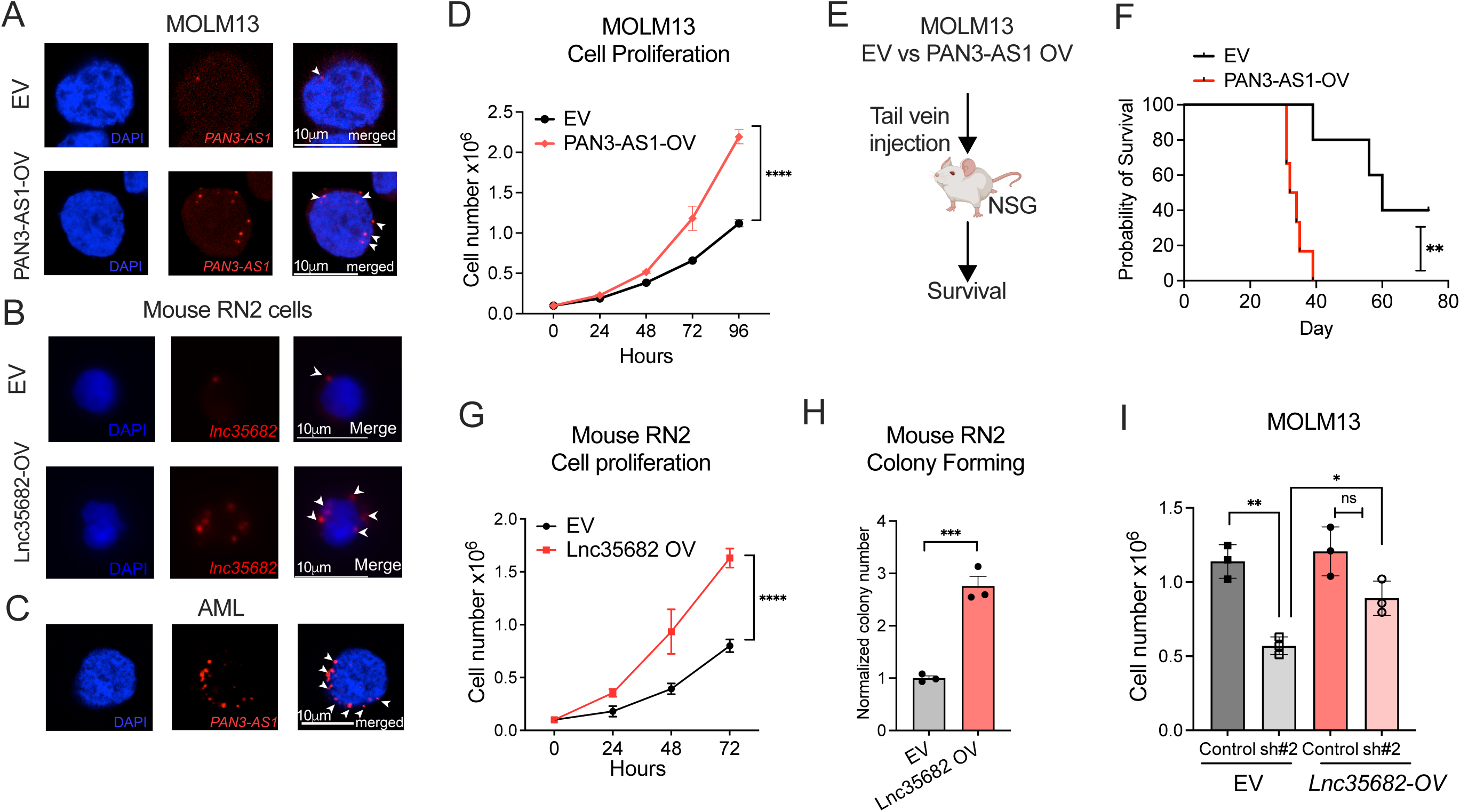
Human *PAN3-AS1* and mouse *Lnc35682* promotes leukemogenesis. (A) *PAN3-AS1* FISH images of MOLM13 cells transduced with lentiviruses carrying an empty vector (EV) or exogenous *PAN3-AS1 c*DNA for overexpression (OV). *PAN3-AS1*: red (highlighted with arrowheads); nucleus: blue (stained with DAPI). Scale bars: 10μm. (B) *Lnc35682* FISH images of mouse RN2 cells transduced with retroviruses carrying an empty vector (EV) or exogenous *Lnc35682* DNA for overexpression (OV). *Lnc35682*: red (highlighted with arrowheads); nucleus: blue (stained with DAPI). Scale bars: 10μm. (C) *PAN3-AS1* FISH images of patient AML cells. *PAN3-AS1*: red (highlighted with arrowheads); nucleus: blue (stained with DAPI). Scale bars: 10μm. (D) Cell proliferation of MOLM13 cells transduced with lentiviruses carrying an empty vector (EV) or exogenous *PAN3-AS1 c*DNA for overexpression (OV). n=3. Data shown, mean ± s.e.m., p-Value: ****<0.0001, two-way ANOVA. (E) Experimental scheme for assessing leukemia progression *in vivo* after transplantation of MOLM13 cells described in (D) (F) Kaplan–Meier survival analysis of mice described in (G). EV (n = 5); OV (n = 6). p-Value: **<0.01, log-rank test. (G and H) RN2 cells transduced with retroviruses carrying an empty vector (EV) or with exogenous *Lnc35682* cDNA for overexpression (OV). (G) Cell proliferation, (H) colony forming assay. n=3. Data shown, mean ± s.e.m., p-Value: ***<0.001, ****<0.0001, two-tailed Student’s t-test. (I) MOLM13 cells with control (scramble) or *PAN3-AS1* KD (sh#2) were transduced with retroviruses carrying an EV or *Lnc35682* DNA for overexpression (OV). *Lnc35682* OV rescued the growth defects of *PAN3-AS1* KD (light red vs light grey). n=3. Data shown, mean ± s.e.m., p-Value: *<0.05, **<0.01, ns, not significant, two-tailed Student’s t-test.

### *PAN3-AS1/Lnc35682* is dispensable for normal hematopoiesis

Next, we sought to explore the roles of *Lnc35682/PAN3-AS1* in normal hematopoietic stem progenitor cells (HSPCs) and determine whether there is a differential requirement for *Lnc35682/PAN3-AS1* function in leukemia cells vs. HSPCs. Knockdown of *Lnc35682* in mouse LSK cells had minimal effects on the colony-forming abilities (**Figure S3A**), indicating that *Lnc35682* is not required for survival and differentiation of normal murine HSPCs *in vitro*. To evaluate its functional requirement *in vivo*, we generated a germline *Lnc35682* knockout (KO) mouse line using CRISPR/Cas9 (**Figure 3A**). *Lnc35682* heterozygous and homozygous KO mice were born at normal mendelian ratio and exhibited no phenotypic defects through their lifetime. Analyses of the hematopoietic system showed equivalent frequencies of HSPC compartment in KO vs. WT littermates (**Figure 3B and S3B**). *Lnc35682* KO and WT bone morrow cells also exhibited comparable repopulating capacity as demonstrated by similar chimerism of CD45.2 positive donor cells in transplantation assay (**Figures 3C-D, and S3B**). In parallel, we transduced LSK cells isolated from KO and WT animals with retroviruses carrying cDNA encoding fusion protein MLL-AF9 prior to injecting the cells into B6 recipient mice (**Figure S3C**). We found that while *Lnc35682* was dispensable for normal hematopoiesis, *Lnc35682* KO delayed MLL-AF9-driven leukemogenesis and lowered disease burdens *in vivo* (**Figures S3D-S3E**).

**Figure 3.**
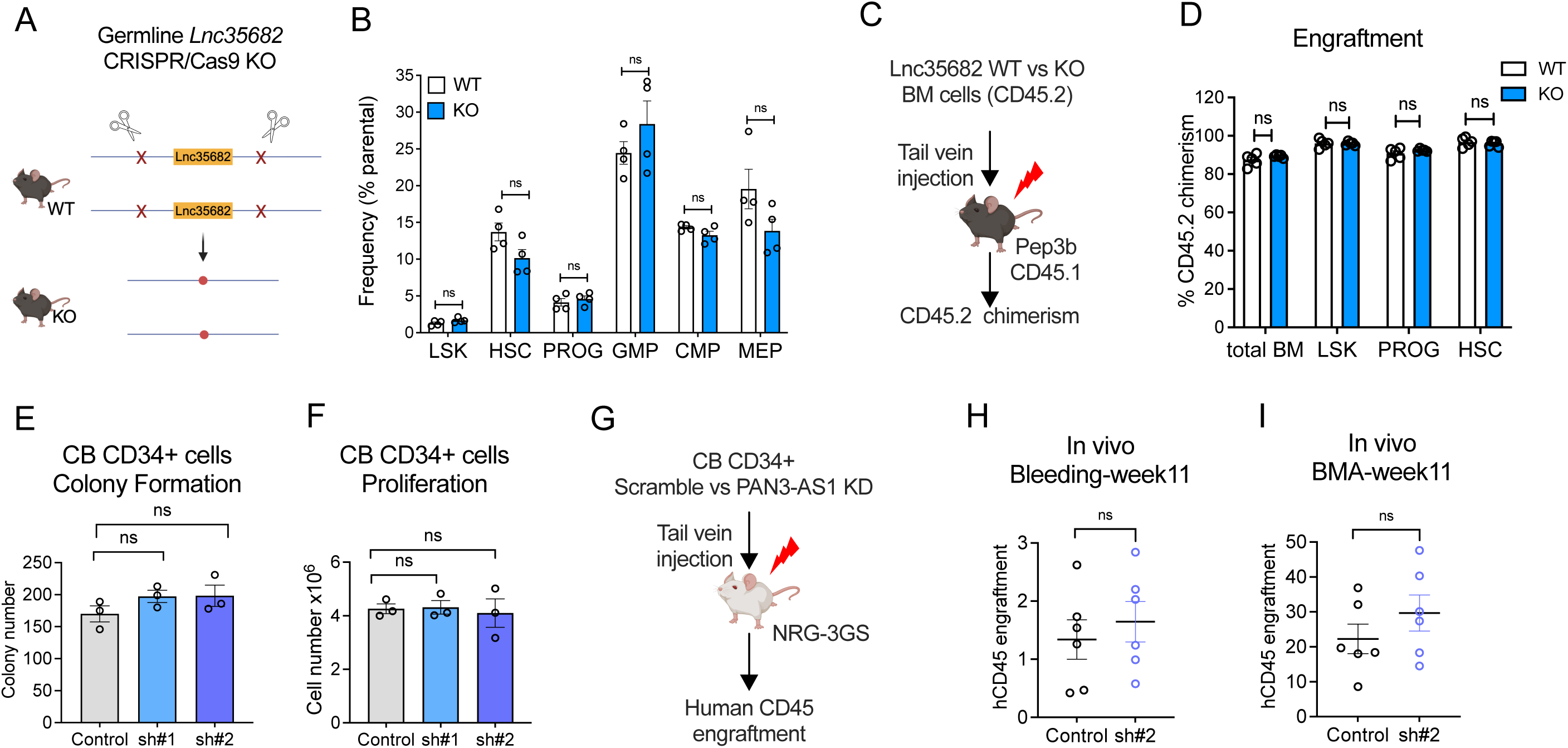
*PAN3-AS1/Lnc35682* is dispensable for normal hematopoiesis. (A) Scheme of generation a CRISPR/Cas9-mediated germline *Lnc35682* knockout (KO, whole gene span homozygous deletion, viable) C57BL/6 mice (B6). (B) Frequencies of stem/progenitor cell compartments acquired from bone marrow (BM) cells of *Lnc35682* WT (hollow bars) and KO (blue bars) littermates. LSK (Lin-Sca+ckit +); HSC – hematopoietic stem cell (LSK CD48-CD150 +); PROG (Lin-Sca-ckit +); CMP – common myeloid progenitor (Lin-Sca+ckit-CD34 + FcγRII/III mid); GMP – granulocyte monocyte progenitor (Lin-Sca+ckit-CD34 + FcγRII/III high); MEP – megakaryocyte erythrocyte progenitor (Lin-Sca+ckit-CD34-FcγRII/III low). n= 4/phenotypes. Data shown, mean ± s.e.m. ns= not significant. (C) Scheme for assessing repopulation potentials of hematopoietic cells from *Lnc35682* wildtype (WT) and homozygous knockout (KO) mice littermates. (D) CD45.2 chimerism in BM of lethally-irradiated recipient mice transplanted with BM from WT and Lnc35682 KO mice littermates at week 16 post-transplantation. N=5/each phenotype. (E and F) Cord blood derived human CD34+ HSPC cells were transduced with lentiviruses expressing either a scramble (control) shRNA or *PAN3-AS1*-targeting shRNAs (sh#1 and sh#2). (E) Colony forming assay, cells were plated on methylcellulose supplemented with cytokines. mean ± s.e.m., Student’s t-test. (F) Cell proliferation in liquid culture supplemented with cytokines. Data shown, mean ± s.e.m., p-Value: ns, not significant, two-tailed Student’s t-test. (G) Experimental scheme for assessing function of normal CD34+ cord blood cell *in vivo* in NRG-3GS mouse. (H) Percentage of human CD45+ cells in peripheral blood (PB) of the recipient NRG-3GS mice described in (G). Control n=6, PAN3-AS1 KD, n=6. Data shown, mean ± s.e.m., p-Value: ns, not significant, two-tailed Student’s t-test. (I) Percentage of human CD45+ cells in bone marrow of the recipient mice described in (G). Control n=6, *PAN3-AS1* KD, n=6. Data shown, mean ± s.e.m., p-Value: ns, not significant, two-tailed Student’s t-test.

In parallel, to directly address the requirement for *PAN3-AS1* in human HSPCs, we performed shRNA-mediated knockdown of *PAN3-AS1* in cord blood (CB) derived CD34+ cells. We observed no reduction in colony formation or proliferation of human HSPCs upon *PAN3-AS1* KD (**Figures 3E-F and S3F-G**). However, overexpression of *PAN3-AS1* resulted in increased colony numbers and enhanced cell growth (**Figures S3H-I**), supporting a leukemia promoting role of *PAN3-AS1.* We further evaluated its functional requirement *in vivo* by transplanting *PAN3-AS1*-KD vs. control CD34+ cells into NRG-3GS mice and followed human cell engraftment (**Figure 3G**). We observed no significant difference in engraftment upon *PAN3-*AS1 depletion in both peripheral blood and bone marrow of recipient mice (**Figures 3H-3I, and S3J**). The data demonstrate that *PAN3-AS1* is not required for normal hematopoiesis, indicating a favourable therapeutic window for suppression of *PAN3-AS1* in AML.

### *PAN3-AS1* functions independently of PAN3

Some antisense lncRNAs have been shown to regulate their sense genes through *cis*-acting mechanisms (Jiang et al., 2023). *PAN3-AS1*, annotated as *PAN3* antisense 1, is transcribed in a head-to-head orientation with *PAN3* on chromosome 13. PAN3 is a subunit of the PAN2/PAN3 RNA deadenylation complex (Christie et al., 2013) and was reported among genes in the Leukemia-Enriched Signature (Mumme et al., 2025). We therefore asked whether *PAN3-AS1* acts via regulation of *PAN3* mRNA expression in AML. Knockdown and overexpression of *PAN3-AS1* resulted in decreased and increased *PAN3* levels, respectively (**Figure S4A-B**), indicating a positive correlation between *PAN3-AS1* and *PAN3* expression. However, we observed that *PAN3* overexpression did not affect *PAN3-AS1* expression (**Figures S4C and S4D**). Importantly, *PAN3* overexpression failed to rescue the growth defects induced by *PAN3-AS1* depletion (**Figure S4E**), indicating that *PAN3-AS1* function is not dependent on *PAN3*. These findings support a non-reciprocal regulatory relationship and suggest that *PAN3-AS1* promotes leukemia through mechanisms extending beyond regulation of its sense gene.

### *PAN3-AS1* influences organization of an oncogenic chromatin state

To unbiasedly identify downstream genes and pathways influenced by *PAN3-AS1* in leukemia cells, we performed transcriptomic profiling following *PAN3-AS1* overexpression and knockdown in MOLM13 cells (**Figure 4A**). Differential expression analysis revealed strong concordance in the genes and pathways altered under these perturbations (**Figures 4B–4G**). Upon *PAN3-AS1* overexpression, 570 genes were upregulated and 323 were downregulated, whereas *PAN3-AS1* knockdown resulted in 861 upregulated and 411 downregulated genes (**Figure 4B-C and *Supplemental Table 3 and 4***). Genes induced by *PAN3-AS1* were enriched for cell proliferation, growth, and developmental pathways, consistent with *PAN3-AS1’s* pro-proliferative role (**Figures 4D-E and *Supplemental Table 5***). Conversely, *PAN3-AS1*–repressed genes were enriched for apoptotic processes, cell differentiation, and inflammatory response pathways, including TNF, IFNγ, and NF-κB signaling (**Figures 4F-G and *Supplemental Table 6***). Notably, transcriptomic profiling further confirmed that *PAN3-AS1* positively regulates *PAN3* expression (**Figures S4F-G**). These data indicate that *PAN3-AS1* promotes a gene expression program supporting leukemia cell growth and survival.

**Figure 4.**
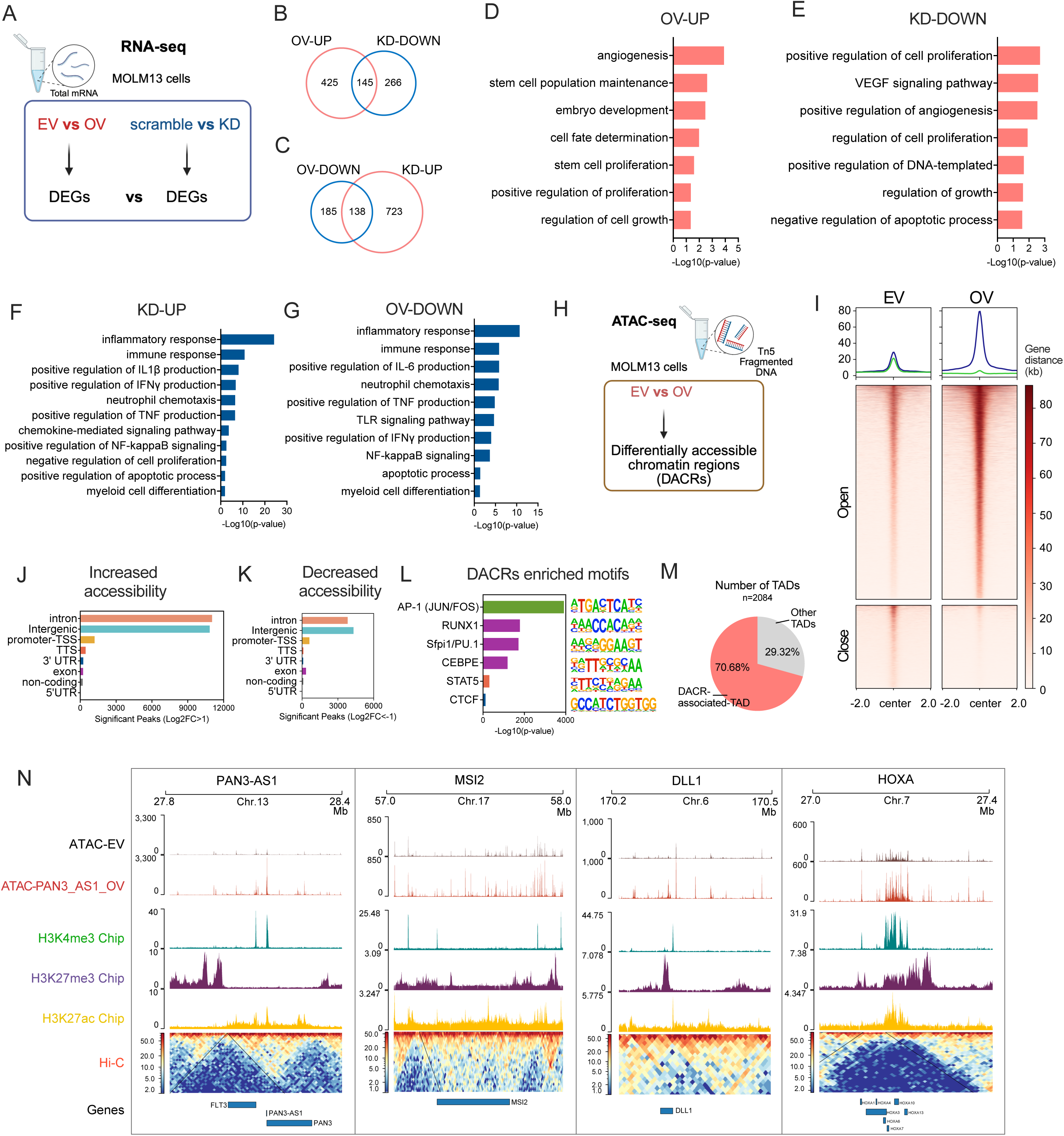
lncRNA *PAN3-AS1*-mediated regulation of a leukemic gene expression program. (A) Scheme of transcriptome profiling (RNA-seq) upon alternation of *PAN3-AS1.* MOLM13 cells were transduced with lentiviruses expressing either scrambled shRNA (control), shRNAs (sh#1, #2) to knockdown (KD) *PAN3-AS1*; or an empty vector (EV) vs. exogenous *PAN3-AS1 c*DNA for its overexpression (OV). Differentially expressed genes (DEGs) were analyzed between groups of *PAN3-AS1*-OV compared to EV, and *PAN3-AS1* KD (sh#1 and sh#2) compared to control. N=3 biological replicates per group. (B-C) Venn diagrams showing numbers of upregulated (UP) and downregulated (DOWN) genes detected in each RNA-seq group and the overlapping numbers of genes with altered expression levels. (D) Gene Ontology Biological Process (GO-BP) analysis of *PAN3-AS1* overexpression (OV) upregulated (UP) genes. (E) GO-BP analysis of *PAN3-AS1* knockdown (KD) downregulated (DOWN) genes. (F) GO-BP analysis of *PAN3-AS1* knockdown (KD) upregulated (UP) genes. (G) GO-BP analysis of *PAN3-AS1* overexpression (OV) downregulated (DOWN) genes. (H) Scheme of ATAC-seq analysis. MOLM13 cells were transduced with lentiviruses carrying an empty vector (EV) vs. exogenous *PAN3-AS1 c*DNA for its overexpression (OV). Differentially accessible chromatin regions (DACRs) were analyzed. N=3 biological replicates per group. (I) Heatmap showing differentially accessible chromatin regions identified by ATAC-seq. Regions showing significant changes in accessibility between *PAN3-AS1* overexpression (OV) and empty vector control (EV) are displayed. Regions are classified as open (log₂FC > 1, p < 0.05, Blue line) or closed (log₂FC < −1, p < 0.05, green line) based on differential accessibility in OV vs. EV samples. (J-K) Genetic feature analysis of the genomic distribution of ATAC-seq differential peaks across promoter-TSS, intron, exon, intergenic, 3’ UTR, 5’ UTR, and non-coding regions in *PAN3-AS1* OV group compared to EV. Regions are classified as Open (log₂FC > 1, p < 0.05) (J) or Closed (log₂FC < −1, p < 0.05) (K) based on differential accessibility in OV versus EV samples. (L) Enriched transcription factor motifs identified in regions with significantly increased chromatin accessibility in OV vs. EV (log₂FC > 1, p<0.05). (M) Pie chart showing the proportion of topologically associating domains (TADs) enriched with differentially accessible chromatin regions (DACRs) in OV vs. EV samples. (N) Genome browser tracks showing representative leukemic gene regions enriched with ATAC-seq signals, together with H3K4me3, H3K27ac and H3K27me3 signals and Hi-C topologically associating domain (TAD) spans the locus.

Given *PAN3-AS1* perinuclear localization, we asked whether *PAN3-AS1* can modulate gene expression by influencing chromatin structure. To explore the hypothesis, we performed ATAC-seq (Assay for Transposase-Accessible Chromatin using sequencing) in *PAN3-AS1* overexpression (OV) cells vs. control (empty vector-EV) to assess chromatin accessibility, which is known to correlate with transcriptional activity (Buenrostro et al., 2013) (**Figure 4H**). We found that *PAN3-AS1* OV led to notable chromatin remodeling, characterized by a substantial gain in accessible regions accompanied by a less extensive loss of accessibility (**Figure 4I**). The differentially accessible chromatin regions (DACRs) were predominantly located within intronic, intergenic and promoter regions, which are genomic compartments enriched for regulatory elements (**Figure 4J-K**). Gene ontology analysis revealed that regions with increased accessibility were significantly associated with genes involved in signaling pathways, cell proliferation, and chromatin remodeling (**Figure S4H and *Supplemental Table 7***), whereas regions with decreased accessibility were linked to pathways including G protein-coupled receptor signal transduction, chemical transport, and cell differentiation (**Figure S4I and *Supplemental Table 8***). Notably, approximately 70% of genes upregulated by *PAN3-AS1* in the RNA-seq analysis overlapped with genes associated with more accessible regions, whereas only 12% of downregulated genes corresponded to regions with decreased accessibility (**Figure S4J**). A number of genes, which exhibited concordance in gene expression changes upon *PAN3-AS1* KD and OV (**Figure 4B-C**), were found to have positive correlation with their decreased and increased accessibility at promoter/transcription start site (TSS) regions (**Figure S4K-L**). Motif enrichment analysis further demonstrated overrepresentation of binding sites of transcription factor families central to hematopoietic and leukemic regulation, including AP-1 (FOS/JUN), RUNX1, PU.1 (Sfpi1), CEBPE, and STAT5, within regions gaining accessibility (**Figure 4L and Supplemental Table 9**). These findings indicate that *PAN3-AS1* promotes leukemia gene expression programs and impacts chromatin accessibility at transcriptional regulatory regions.

Given that regulatory regions are usually organized into higher-order chromatin structures, we examined whether the chromatin accessibility gains upon *PAN3-AS1* overexpression are embedded within chromatin compartments. We integrated our ATAC-seq data with publicly available Hi-C and histone ChIP-seq datasets generated for leukemia cells (Klco et al., 2023; Xu et al., 2022). Notably, we observed that over 70% of DACRs - ATAC peaks aligns with topologically associating domains (TADs) defined by Hi-C mapping (Xu *et al*., 2022) (**Figure 4M**). The majority of genomic regions that interact with the *PAN3-AS1* locus displayed changes in chromatin accessibility (**Figure S4M**). In addition, we noticed overlaps of regions with increased accessibility with enrichments of active histone marks (H3K4me3 and H3K27ac) and depletion of the repressive mark H3K27me3, notably within regulatory regions of critical leukemia loci such as *MSI2*, *DLL1* and *HOXA* (**Figure 4N**). Together, these findings supported a model in which elevated *PAN3-AS1* expression can impact chromatin structure preferentially within spatially proximal chromatin regulatory hubs. As a result, *PAN3-AS1* engages with a *trans*-regulatory program that facilitates activation of leukemic gene expression programs to drive disease progression.

### *PAN3-AS1* engagement with Lamin B1 is essential for promoting leukemia cell growth

Given the observed subcellular localization of *PAN3-AS1* and its effects on chromatin accessibility, we asked whether *PAN3-AS1* associates with the nuclear lamina to modulate chromatin organization. We chose to assess *PAN3-AS1* potential interaction with Nuclear lamin B1 (LMNB1), a key component of the nuclear envelope, which anchors chromatin to the nuclear periphery through lamina-associated domains and is involved in 3D chromosomal organization (Camps et al., 2015; Reilly et al., 2022; Shin et al., 2026). *PAN3-AS1* RNA FISH combined with LMNB1 immunofluorescence staining demonstrated co-localization of *PAN3-AS1* and LMNB1 around the perinuclear region (**Figures 5A and S5A**). To further confirm *PAN3-AS1* and LMNB1 association, we performed LMNB1 RNA Immunoprecipitation quantitative PCR (RIP-qPCR) and found significant *PAN3-AS1* enrichment upon LMNB1 immunoprecipitation (**Figures 5B-C**). These data suggest a functional interaction between the nuclear lamina and *PAN3-AS1* in AML cells.

**Figure 5.**
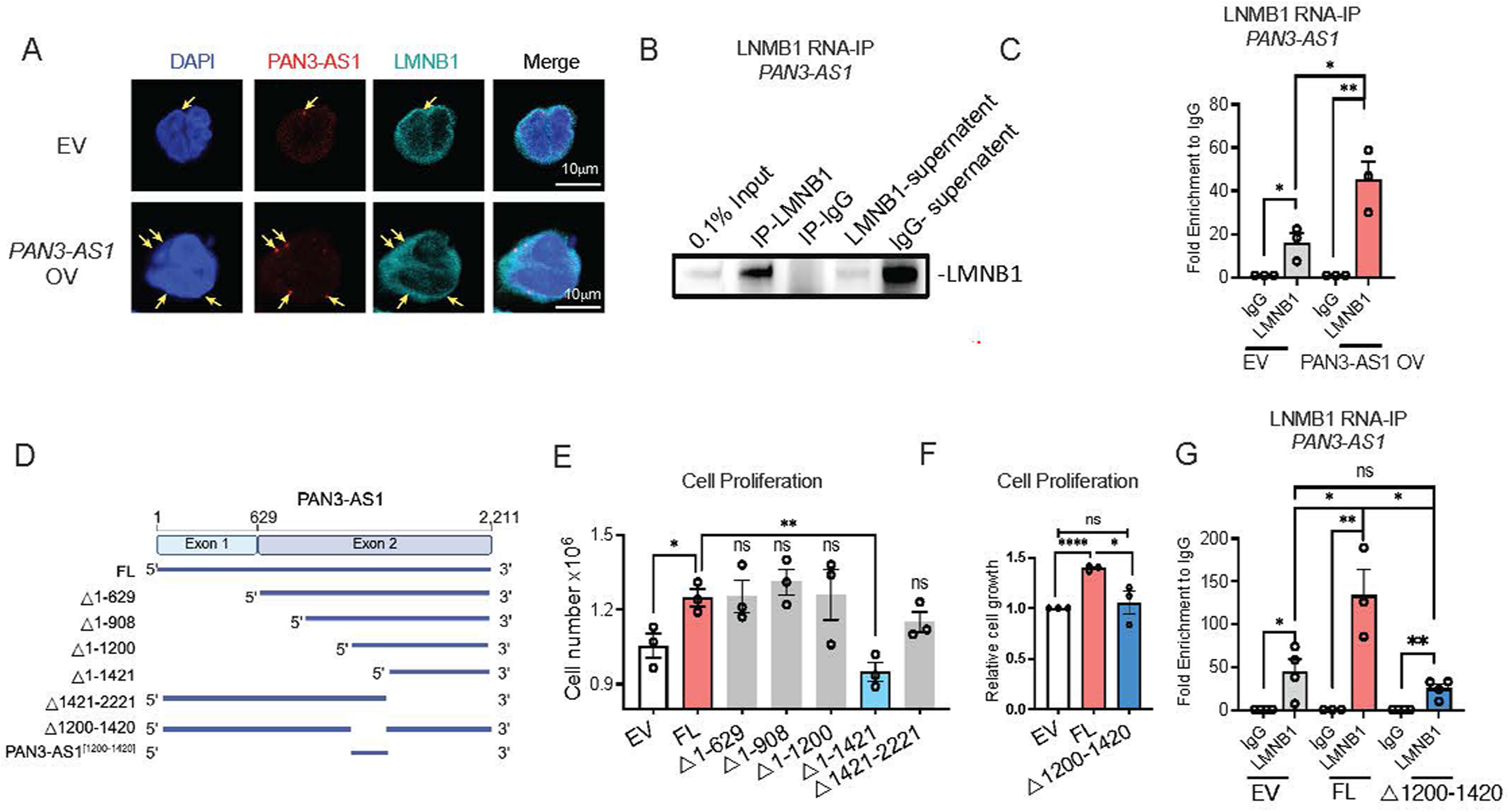
*PAN3-AS1* associates with Lamin B1 via a conserved region. (A) RNA FISH and LNMB1 immunofluorescence analyses of MOLM13 cells transduced with empty vector (EV) or *PAN3-AS1* overexpression construct (OV). *PAN3-AS1* RNA signals (red, arrows) co-localize with LMNB1 (cyan) in perinuclear regions. Nuclei were stained with DAPI (blue). Scale bars: 10μm. (B) Immunoblot confirming efficient immunoprecipitation of LMNB1 in the RNA-IP assay. (C) RIP–qPCR analysis showing fold enrichment of *PAN3-AS1* following LMNB1 immunoprecipitation relative to IgG control in EV or *PAN3-AS1* OV cells. (D) Schematic of full length (FL) and truncated *PAN3-AS1* fragments used for functional analysis. (E) Cell proliferation of MOLM13 cells transduced with EV or constructs expressing different *PAN3-AS1* fragments. (F) Cell proliferation of MOLM13 cells transduced with EV, constructs expressing full length *PAN3-AS1* (FL), or with deletion of [1200-1420] (△[1200-1420]). (G) RIP–qPCR analysis showing fold enrichment of *PAN3-AS1* following LMNB1 immunoprecipitation relative to IgG control in EV, FL, and △[1200-1420] cells.

Next, we reasoned that if *PAN3-AS1*’s association with LMNB1 is key to its leukemia promoting role, the interaction is likely mediated by a conserved functional region within the lncRNA. Thus, to define the regions of *PAN3-AS1* underlying its function and interaction with LMNB1, we first looked into conservation by cross-species sequence alignment (Gaonac’h-Lovejoy et al., 2025). The analysis revealed that exon 1 was less conserved across species than exon 2 (**Figures S5B**). To then experimentally map the minimally functional domains of *PAN3-AS1*, we generated a series of truncation mutants, overexpressed them in MOLM13 cells and evaluated their effects on AML cell growth (**Figure 5D**). We observed that the 5’ region [1-1200] and 3’ region [1421-2221] were dispensable for *PAN3-AS1*’s growth induction while removal of the sequence PAN3-AS1△[1-1421] markedly attenuated the growth-promoting effect of *PAN3-AS1* (**Figures 5E),** indicating that the more conserved middle region [1200-1420] (**Figure S5C)** might be required to *PAN3-AS1* activity. Indeed, deletion of the internal [1200-1420] segment was sufficient to abrogate *PAN3-AS1* positive effect on AML cell proliferation (**Figure 5F**) and overexpression of the fragment alone resulted in enhanced cell growth (**Figures S5D and S5E**). Moreover, LMNB1 RIP-qPCR analyses demonstrated that loss of PAN3-AS1△[1200-1420] diminished *PAN3-AS1* binding to LMNB1, indicating that this conserved domain is required for *PAN3-AS1*-LMNB1 interaction (**Figure 5G**). These data suggest that elevated *PAN3-AS1* expression in leukemia cells enhances its interaction with LMNB1, thus influencing chromatin accessibility that favor the expression of leukemia-promoting genes and ultimately promote leukemogenesis.

### *PAN3-AS1* and FLT3 form a *cis*-regulatory module

Having established the importance of *PAN3-AS1* across several experimental models of AML, we next sought to assess its clinical relevance. We surveyed *PAN3-AS1* expression in AML subtypes within the 4 major patient cohorts i.e., Beat AML (Bottomly et al., 2022), TCGA (Cancer Genome Atlas Research et al., 2013), Leucegene (Lavallée et al., 2016) and BC Cancer AML Personalized Medicine Project (AML-PMP) (Docking et al., 2021). We saw that *PAN3-AS1* expression was elevated in high-risk and normal karyotype AML subtypes, specifically those harboring FLT3-ITD, NPM1, and IDH1/2 mutations (Papaemmanuil et al., 2016) (**Figures 6A, S6A-B**). We were particularly interested in FLT3 mutations, which are among the most common mutations in AML and account for approximately 25% of all AML cases (Daver et al., 2019; Papaemmanuil *et al*., 2016). *FLT3* gene is located in the same conserved genomic region as *PAN3-AS1* on chromosome 13. Independent analysis using the Alliance AML patient cohort (Giacopelli et al., 2021) confirmed higher *PAN3-AS1* expression in genetic subtypes of FLT3, NPM1 and IDH2 mutations (data not shown). Interestingly, within the Alliance cohort*, PAN3-AS1* expression was found to exhibit a stronger correlation with *FLT3* than with *PAN3*, despite *PAN3-AS1* and *PAN3* sharing the same promoter (**Figure S6C**). We also observed that the presence of the STAT Hypomethylation Signature (SHS), which captures AML epitypes exhibiting STAT-associated hypo-DNA methylation regardless of FLT3 mutation status, stratified *PAN3-AS1* and *FLT3* high expressing patients (Giacopelli *et al*., 2021), suggesting a connection between activation of FLT3 signaling pathway and upregulation of *PAN3-AS1* and *FLT3* expression in AML patients (**Figures S6D**). On the other hand, differential gene expression analysis comparing *PAN3-AS1*^high^ versus *PAN3-AS1*^low^ patient groups in all 4 Beat AML, TCGA, Leucegene and AML-PMP cohorts revealed higher expression of *FLT3* in PAN3-AS1^high^ patient group (**Figures S6E).** These data suggest a potential link between FLT3 activation and *PAN3-AS1* upregulation in AML.

**Figure 6.**
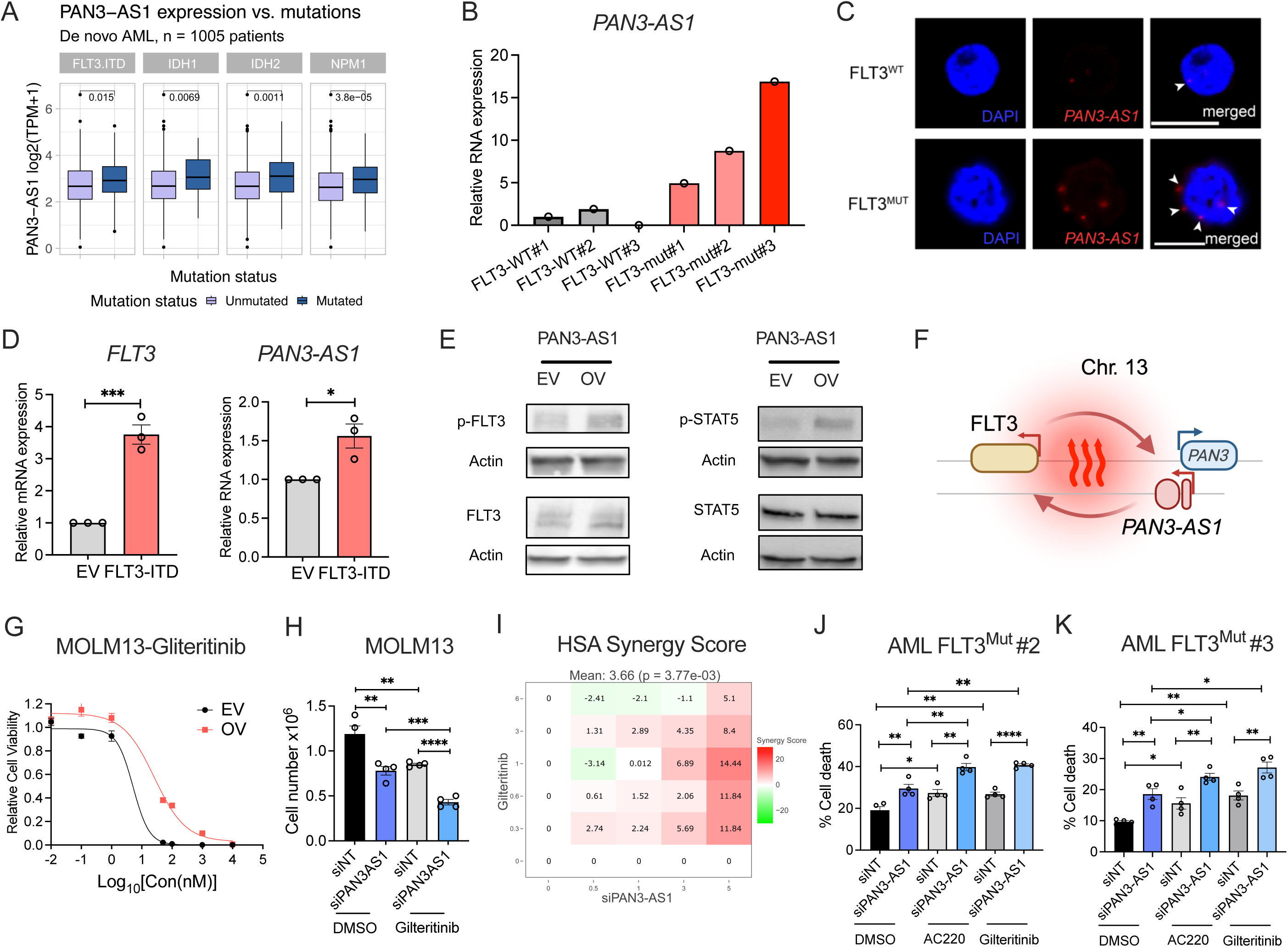
*PAN3-AS1* and FLT3 forms a cis-regulatory module in leukemia. (A) *PAN3-AS1* expression levels in *de novo* AML patient cohorts categorized based on mutation status within the cytogenetic normal karyotype AML (BEAT-AML, Leucegene, PMP, and TCGA). (B) Quantitative RT-qPCR analysis of *PAN3-AS1* expression levels in FLT3 wildtype (FLT3^WT^) and FLT3 mutant (FLT3^mut^) primary patient AML cells. (C) *PAN3-AS1* FISH RNA scope images of FLT3 wildtype (FLT3^WT^) and FLT3 mutant (FLT3^mut^) patient AML cells. More abundant *PAN3-AS1* signals (red, arrowheads) were observed in perinuclear regions (nucleus marked by DAPI, blue) in the FLT3^mut^ than FLT3^WT^ cells. Scale bars: 10μm. (D) Quantitative RT-qPCR analysis of *FLT3* (left) and *PAN3-AS1* (right) expression levels in THP-1 (an AML cell line without harboring FLT3 mutation) transduced with of empty vector (EV) or overexpressing exogenous *FLT3-ITD* cDNA. n=3. Data shown, mean ± s.e.m., p-Value: *<0.05, ***<0.001, two-tailed Student’s t-test. (E) Immunoblots showing FLT3, phospho-FLT3 (pFLT3), STAT5, phospho-STAT5 (pSTAT5) protein expression upon *PAN3-AS1* overexpression vs. EV control. ACTIN serves as loading control. (F) Schematic of the proposed FLT3 and PAN3-AS1 functional interacting loop. (G) Dose-response curves showing effects of FLT3 inhibitor Gilteritinib on MOLM13 cells expressing empty vector (EV) vs. *PAN3-AS1* overexpression cDNA (OV). Relative cell viability was measured by CellTiter-Glo at 72 h after drug treatment (n = 3). (H) Effects of Gilteritinib (1nM) on MOLM13 cells treated with control siRNA (siNT) or *PAN3-AS1*– targeting siRNA (siPAN3-AS1). Cell numbers were counted at 72 h after drug treatment. n=4. Data shown, mean ± s.e.m., p-Value: **<0.01, ***<0.001, ****<0.0001, two-tailed Student’s t-test. (I) Synergy score of Gilteritinib treatment combined with *PAN3-AS1* depletion in MV4-11 cells, calculated using SynergyFinder. (J and K) Effects of Gilteritinib (1nM) and Quizatinib (AC220, 0.5nM) on primary patient FLT3^mut^ AML cells (#2 in J, #3 in K) treated with control siRNA (siNT) or *PAN3-AS1*–targeting siRNA (siPAN3-AS1). Percentages of trypan blude stained cell death were scored at 72 h after treatment. n=4. Data shown, mean ± s.e.m., p-Value: *<0.05, **<0.01, ****<0.0001, two-tailed Student’s t-test.

Given the clinical observations, we confirmed elevated levels of *PAN3-AS1* in FLT3 mutant AML by comparing *PAN3-AS1* expression levels in our in-house AML patients carrying FLT3 mutations (FLT3^mut^) or FLT3 wild type (FLT3^WT^) (**Figure 6B-C**). To address whether activation of the FLT3 signaling pathway in FLT3^mut^ AML indeed results in upregulation of *PAN3-AS1*, we ectopically expressed the FLT3 Internal Tandem Duplication (ITD) mutant- *FLT3-ITD* in non-FLT3 mutation THP-1 leukemia cells. We found that *FLT3-ITD* could significantly increase *PAN3-AS1* expression (**Figure 6D**). Conversely, we demonstrated that *PAN3-AS1* depletion and overexpression resulted in reduction and increase *FLT3* expression, respectively (**Figures S6F-G**). *PAN3-AS1* overexpression led to higher levels of FLT3 protein and phosphorylated FLT3 (**Figure 6E and S6H-I**). In addition, we observed an increase in phosphorylation of STAT5, the key downstream effector of the FLT3-STAT5 signaling axis (**Figures 6E and S6J-K**). Constitutive FLT3–STAT5 signaling is a key oncogenic pathway in FLT3 mutant AML, driving leukemia proliferation and survival (Spiekermann K, 2003). Our findings suggest that acquisition of FLT3 mutations can lead to *PAN3-AS1* upregulation, which in turn remodel chromatin accessibility to further promote *FLT3* expression and enhance leukemic gene expression programs. Collectively, these results demonstrated a functional feedforward loop between FLT3 and *PAN3-AS1* driving leukemia progression (**Figure 6F**). Considering the closer association between *PAN3-AS1* and FLT3 signaling, but not the *PAN3* sense gene, we, therefore, proposed describing *PAN3-AS1* as lncRNA *FLAMER* (<u>F</u>LT3-<u>L</u>inked <u>AM</u>L <u>E</u>nhancer-like <u>R</u>NA) to better capture this aspect of its biological function.

Given the use of FDA approved FLT3 inhibitors AC220 (Quizatinib) and Gilteritinib to treat FLT3 mutant AML in clinics (Cortes et al., 2019; Erba et al., 2023; Kennedy and Smith, 2020; Perl et al., 2017; Perl et al., 2019), we next sought to evaluate whether *PAN3-AS1* activity influences the sensitivity of AML to FLT3 inhibition. We treated *PAN3-AS1* overexpression vs. EV control MOLM13 cells with a range of concentrations of AC220 and Gilteritinib and tracked cell viability at 72h after treatment. We observed that *PAN3-AS1* high expression conferred strong resistance to both of the drugs (**Figures 6G and S6L**). Conversely, knocking down of *PAN3-AS1* using either shRNA or siRNA potentiated the killing effects of FLT3 inhibitors with considerable synergistic effects in MOLM13 and MV4-11 leukemia cells harboring FLT3-ITD mutations (**Figures 6H, 6I, S6M-O**). Consistent with our findings in cell lines, the combination treatment exhibited enhanced efficacy in primary FLT3-mutant AML patient cells (**Figures 6J-K, S6P-Q**). Taken together, these data provide a pre-clinical rationale for therapeutic targeting of *PAN3-AS1* in FLT3 mutant leukemia.

### Development of a Lipid nanoparticle (LNP) - based siRNA therapy for myeloid leukemia

Lipid nanoparticle (LNP)–mediated RNA delivery has recently emerged as a promising platform for nucleic acid therapeutics(Cullis and Felgner, 2024). However, conventional LNP formulations predominantly accumulate in the liver, exhibit short circulation time, and limited delivery to hematopoietic tissues(Hosseini-Kharat et al., 2025). To evaluate the feasibility of LNP-mediated siRNA delivery into leukemia cells, we tested a newly developed LNP formulation, LNP^HiPEH^ (high potency extrahepatic lipid nanoparticle), incorporating several modifications to the lipid composition compared to the clinically available siRNA-LNP formulation Patisiran (**Figure 7A**). The ionizable lipid content was reduced, while the proportion of helper lipids was increased to promote formation of liposomal LNP structures with prolonged circulation and enhanced extrahepatic delivery (Cheng et al., 2025). In addition, we boosted buffer ionic strength during formulation to induce structural features that improve RNA stability and transfection efficiency (Cheng et al., 2023). We also optimized the sodium citrate concentration to preserve RNA loading while enabling improved delivery to hematopoietic organs, including the spleen and bone marrow (*manuscript is under review*). DiI (1,1’-dioctadecyl-3,3,3’,3’ tetramethyl-lindocarbo-cyanine perchlorate), a lipophilic membrane dye was added alongside the other lipids during in LNP formulation, allowing detection of LNP uptake in cells by flow cytometry analysis. LNP^HiPEH^ displayed minimal toxicity in a wide range of dosages (**Figure S7A**) and nearly 100% uptake in AML cells (**Figure S7B**). To examine the cell-type specificity of LNP delivery, we also assessed LNP^HiPEH^ uptake in normal hematopoietic cells. We noted that human CB CD34+ HSPC cells showed highest LNP^HiPEH^ uptake among various populations of hematopoietic cells (**Figure S7C-E**). In general, compared to AML, the majority of normal human and mouse hematopoietic cells showed relatively lower level of LNP^HiPEH^ uptake (**Figures S7C-E**).

**Figure 7.**
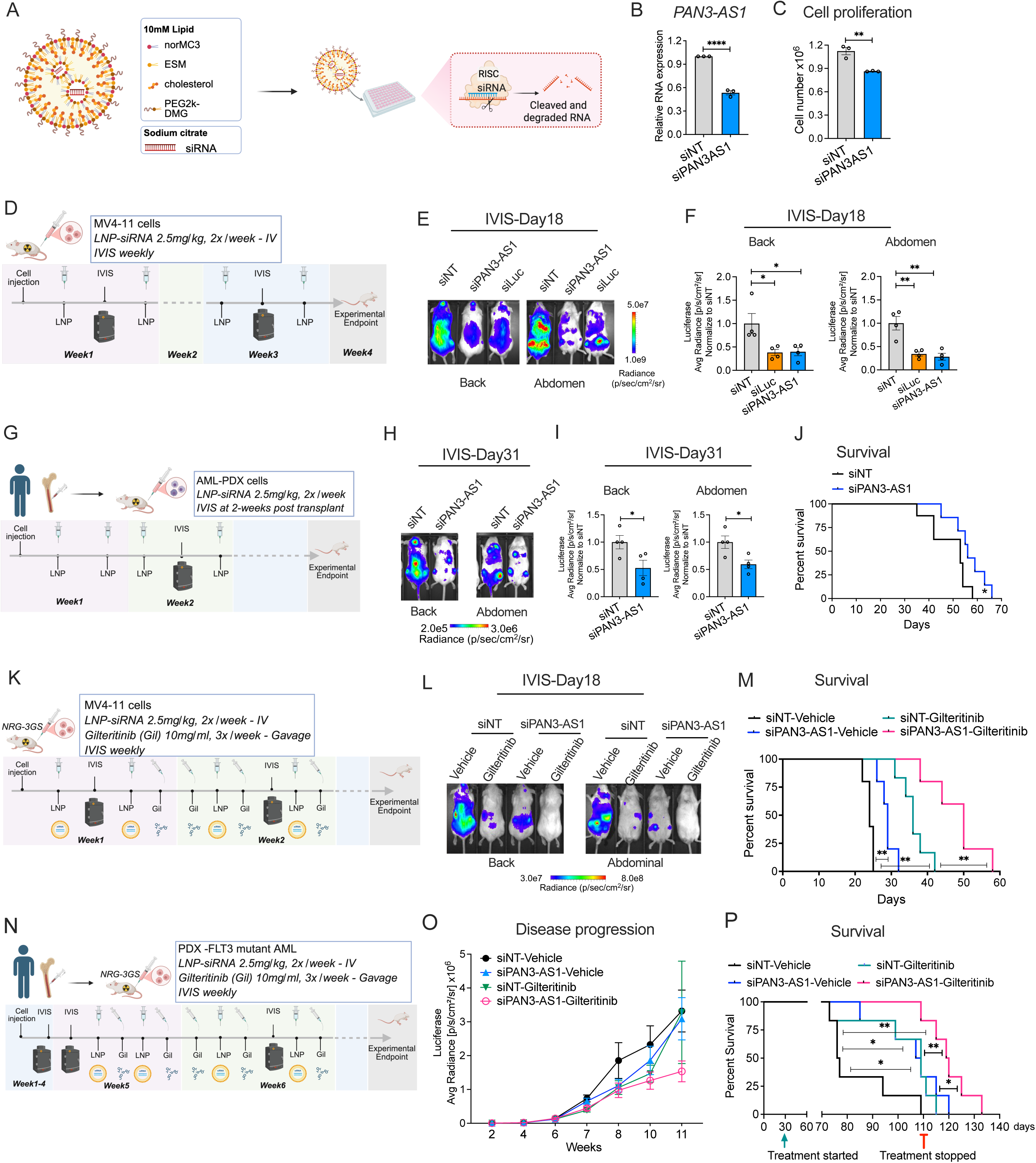
siRNA packed LNP enabled effective *PAN3-AS1* targeting for AML treatment. (A) Schematic of the design of the AML–targeting LNP formulation (LNP^HiPEH^) and LNP–siRNA delivery to deplete RNA in AML cells. (B) Quantitative RT-qPCR analysis *of PAN3-AS1* expression levels in MV4-11 cells upon 48h post-transfection with LNP^HiPEH^-siPAN3-AS1 (si#3) vs. non-targeting siRNA (siNT). (C) Reduced MV4-11cell growth at 72h post transfection described in (B). (D) Experimental scheme for *in vivo* assessment of leukemia progression. MV4-11 cells were engineered to express CBR-Luciferase, which allows for tracking of leukemia burden in animals by IVIS imaging. 8×10^4^ MV4-11 cells were transplanted into NRG-3GS mice prior to receiving treatments with LNP^HiPEH^-siPAN3-AS1, LNP^HiPEH^-si-CBR-Luciferase (siLuc), or LNP^HiPEH^-siNT. (E) IVIS imaging at day 18 post xenograft of MV4-11 cells in NRG-3GS mice described in (D). (F) Quantification of luminescent signals from the dorsal (back) and abdominal sides (abdomen) of mice shown in (E) (n = 4). Data are presented as mean ± s.e.m., p-Value: *< 0.05, **< 0.01, two-tailed Student’s t-test. (G) Experimental scheme for *in vivo* assessment of PDX leukemia progression. PDX cells were engineered to express Luciferase, which allows for tracking of leukemia burden in animals by IVIS imaging. 3×10^5^ PDX cells were transplanted into NRG-3GS mice prior to receiving treatments with LNP^HiPEH^-siPAN3-AS1 or siNT. (H) IVIS imaging at day 31 post xenograft in PDX cells in NRG-3GS mice described in (G). (I) Quantification of luminescent signals from the dorsal (back) and abdominal sides (abdomen) of mice shown in (H) (n = 4). Data are presented as mean ± s.e.m., p-Value: *< 0.05, two-tailed Student’s t-test. (J) Kaplan–Meier survival analysis of mice described in (G). siNT, n = 8; siPAN3-AS1, n = 7. p-Value: *<0.05, log-rank test. (K) Experimental scheme for *in vivo* assessment of PDX leukemia progression. Luciferase MV4-11 cells were transplanted into NRG-3GS mice prior to receiving treatments with vehicle, Gilteritinib, LNP^HiPEH^-siPAN3-AS1, or siNT. (L) IVIS imaging at day 18 post xenograft in NRG-3GS mice described in (K). (M) Kaplan–Meier survival analysis of mice described in (K). n=6 for the siNT-Gilteritinib group, N=5 for the other three groups. is shown in the graph. p-Value: **<0.01, log-rank test. (N) Experimental scheme for *in vivo* assessment of AML-PDX leukemia progression. FLT3^mut^ (#3) PDX cells were engineered to express CBR-luciferase, which allows for tracking of leukemia burden in animals by IVIS imaging. 3×10^5^ PDX cells were transplanted into NRG-3GS mice expansion for 4 weeks prior to receiving treatments with vehicle, Gilteritinib, LNP^HiPEH^-siPAN3-AS1, or siNT. (O) IVIS-based longitudinal monitoring of leukemia burden in the mice described in (N). (P) Kaplan–Meier survival analysis of mice described in (N). n=6. p-Value: *<0.05; **<0.01, log-rank test. Treatment started and stopped points were indicated in the graph.

Next, we injected LNP^HiPEH^ to evaluate LNP uptake *in vivo*. We observed that while LNP uptakes by hematopoietic cells in bone marrow, liver and spleen were minimal (**Figure S7F**), LNP targeting to leukemia cells was very efficient (**Figure S7G**). In parallel, we optimized silencing of *PAN3-AS1* in MOLM13 leukemia cells by testing 3 independent siRNAs to select siRNA#3 (named siPAN3-AS1 hereafter), which showed strongest inhibition against *PAN3-AS1* for subsequent LNP^HiPEH^ packaging and delivery (**Figures S7H-J**). *In vitro* transfection of LNP^HiPEH^ - siPAN3-AS1 in MOLM13 and MV4-11 cells resulted in efficient knockdown of *PAN3-AS1* expression (**Figures 7B and S7J**) and inhibition of *in vitro* cell growth compared to non-targeting (siNT) control (**Figure 7C**). Altogether, these results were particularly encouraging for the potential *in vivo* delivery of LNP siPAN3-AS1 payloads.

### *In vivo* treatment of AML with LNP-siPAN3-AS1

Next, we evaluated the efficacy of LNP^HiPEH^ -siPAN3-AS1 therapy in AML xenograft models. To enable real time tracking of *in vivo* disease progression, we engineered leukemia cells to express Click Beetle Red-Luciferase (CBR-LUC), which allows for measurement of leukemia burden in animals with the IVIS (In Vivo Imaging System). To orthogonally validate on-target LNP delivery, we obtained siRNA targeting CBR-LUC (siLuc), which demonstrated effective gene knockdown and reduction of luciferase signals upon delivery via LNP^HiPEH^ (**Figure S7K-L**). We first tested the feasibility and efficacy of siRNA silencing of endogenous targeting with LNP^HiPEH^ delivery in our well-established MV4-11 leukemia xenograft model (**Figure 7D**). We injected 8×10^4^ MV4-11 expressing luciferase (MV4-11 LUC) into sublethally irradiated NRG-3GS mice. 24h after cell injection, LNP^HiPEH^, packed with siNT, siLuc, and siPAN3-AS1, were administrated intravenously (IV) twice a week at the 2.5mg/kg dosage. We did not observe any overt adverse effects on the general health of treated animals, indicating that the LNP treatment was well tolerated. To track leukemia burden and progression, we performed IVIS once a week until the majority of mice succumbed to the disease. We observed reduced luciferase levels in mice received either LNP^HiPEH^ -siLuc and LNP^HiPEH^-siPAN3-AS1 in comparison to those treated with LNP^HiPEH^-siNT (**Figures 7E-F, S7M-N**). While decrease in luciferase signal in LNP^HiPEH^-siPAN3-AS1 treated mice reflected reduced leukemia burden (**Figure S7O-P**), reduction of luciferase signals proved that siRNA targeting was on-target and independent of disease states. Thus, the data strongly indicated that siRNAs were effectively delivered and silenced target genes to leukemia cells. Subsequently, to demonstrate the relevance of the approach in primary diseases, we applied the LNP^HiPEH^-siPAN3-AS1 treatment regimen in a patient derived xenograft (PDX) model (**Figure 7G**). Consistent with the MV4-11 cell line model, LNP^HiPEH^-siPAN3-AS1 led to lower disease burden (**Figures 7H-I, S7Q-S)** and improved survival (**Figures 7J**). We noted that the expanded leukemia cells collected from the endpoint of LNP^HiPEH^-siPAN3-AS1 treated mice showed comparable *PAN3-AS1* expression with those treated with LNP^HiPEH^-siNT, suggesting that *PAN3-AS1* depleted cells were negatively selected against *in vivo* and leukemia progressed under LNP^HiPEH^-siPAN3-AS1 treatment were driven by cells successfully evaded RNAi mediated gene silencing (**Figure S7T**). Altogether, these results demonstrate that LNP^HiPEH^-siPAN3-AS1 could serve as a new strategy for treatment of AML.

### Combination therapy targeting the FLT3-*PAN3-AS1* regulatory loop is effective in AML

Given the combinatorial effects of FLT3 inhibition and *PAN3-AS1* silencing, as well as the demonstrated feasibility of *in vivo* siPAN3-AS1 delivery, we sought to determine whether combining FLT3 inhibitors with LNP^HiPEH^-siPAN3-AS1 could yield improved therapeutic efficacy in FLT3-mutant AML. Using the FLT3-ITD mutant MV4-11 LUC xenograft model, we set up 4 treatment arms where recipient NRG-3GS mice were treated with (i) LNP^HiPEH^-siNT and vehicle; (ii) LNP^HiPEH^-siNT and Gilteritinib; (iii) LNP^HiPEH^-siPAN3-AS1 and vehicle; and (iv) LNP^HiPEH^-siPAN3-AS1 and Gilteritinib. We tracked disease progression by IVIS weekly and followed overall survival of treated animals (**Figure 7K**). Treatment with FLT3 inhibition or *PAN3-AS1* silencing alone, as well as the combination treatment, resulted in a marked reduction in leukemia burden at Day 11 (**Figure S7U)** and Day 18 checkpoints (**Figures 7L and S7V)**. Notably, combination treatment significant hindered disease progression and improved survival of treated animals (**Figure 7M**).

To evaluate the applicability of our findings in a more clinically relevant setting, we established a transplantable FLT3-mutant AML PDX model, engineered the PDX cells to express luciferase, and administered the four treatment regimens described above (**Figure 7N**). Given the longer disease kinetic of this model and closely mimicked treatment in patients with an established disease, we confirmed leukemia engraftment at week 4 post transplantation of the PDX cells prior to starting the treatment. We observed that while the leukemia inhibitory effects were comparable across treatment groups at early time points, combination of Gilteritinib and LNP^HiPEH^-siPAN3-AS1, in comparison to single arm treatments with either Gilteritinib or LNP^HiPEH^-siPAN3-AS1, led to more sustainable suppression of disease progression beyond week 8 check point (**Figure 7O**). Importantly, Gilteritinib and LNP^HiPEH^-siPAN3-AS1 combination treatment resulted in significantly better overall survival of experimental animals (**Figure 7P**). Together, our study provides strong preclinical evidence supporting a novel therapeutic strategy that exploits a targetable vulnerability through the combination of FLT3 inhibitors and LNP^HiPEH^-siPAN3-AS1 in AML.

## Discussion

Long non-coding RNAs have emerged as important regulators of leukemia biology, yet only a limited number have been functionally characterized, and even fewer have been evaluated as therapeutic targets in clinically relevant models. While previous studies identified several oncogenic or tumor suppressor lncRNAs in leukemia context, their translational significance has often been constrained by limited conservation between species, incomplete validation in primary patient samples and the absence of effective in vivo targeting strategies (Lyu *et al*., 2017; Zeng *et al*., 2014; Zhang *et al*., 2018; Zhang *et al*., 2016). In addition, the therapeutic window associated with targeting many leukemia-relevant lncRNAs has not been established. Our study addresses several of these limitations by identifying *PAN3-AS1* as a therapeutically actionable lncRNA that is preferentially expressed and functionally required in AML. Using complementary genetic and molecular approaches, we demonstrate that *PAN3-AS1* supports leukemia cell survival while showing limited requirement in normal hematopoietic cells, displaying a favourable therapeutic window. Importantly, we demonstrated its functional importance across distinct *in vivo* AML models, including genetic mouse models, cell line xenografts and several clinically relevant PDXs. Notably, the results showing effective depletion of *PAN3-AS1* and suppression of leukemogenesis using systemically delivered siRNA-loaded LNPs further distinguishes our study from earlier work that primarily demonstrated biological function without establishing a feasible therapeutic strategy. Together, our work provides solid evidence that lncRNA dependencies can be identified, mechanistically characterized, and therapeutically exploited in AML.

Our findings support a model in which *PAN3-AS1* contributes to leukemia-promoting gene regulation through both local and broader chromatin states. *PAN3-AS1* is positioned adjacent to *FLT3* and appears to participate in a *cis*-regulatory module that promotes *FLT3* expression and signalling while itself is impacted by activation of the locus under influence of leukemia associated mutations in FLT3 and other epigenetic regulators e.g., IDH1/2 and NPM1. The chromatin-interaction and accessibility analyses showed that *PAN3-AS1* may influence multiple genomic regions brought into spatial proximity with its locus. These results suggest that *PAN3-AS1* upregulation in AML patients is likely a result of aberrant chromatin regulation initiated by these founding mutations. The increase in the *PAN3-AS1* lncRNA abundance, through affecting LNMB1 interactions within impacted genomic regions, then feeds into the regulatory loop, further sustains transcription programs that drive disease progression (**graphical abstract**). Our data pinpointing the critical functional domain required for *PAN3-AS1* leukemia promoting activity and its association with LNMB1 also corroborates the proposed mechanism. At the same time, while we presented several lines of evidence supporting this model, it still remains to be determined whether *PAN3-AS1* directly anchors specific genomic regions to the nuclear lamina or instead modulates chromatin architecture through a more dynamic or indirect mechanism. Given the ubiquitous distribution of lamin proteins around the nuclear periphery, we favor a scenario in which *PAN3-AS1* associates with local LMNB1 protein, thereby rearranging spatially proximal genomic regions within distinct lamina-associated domains (LADs), possibly repelling interactions of some gene loci from a suppressive chromatin environment. Emerging evidence underscores the dynamic and heterogeneous nature of LADs (Briand and Collas, 2020), the existence of facultative LADs alongside constitutive LADs (Alagna et al., 2023; Yanez-Cuna and van Steensel, 2017), suggesting a more complex relationship between LAD organization and transcriptional activity. In addition, the widespread increase in chromatin accessibility following *PAN3-AS1* upregulation may partly reflect secondary effects mediated through enhanced activity of master transcription factors, including AP-1, RUNX1, PU.1, CEBP, and STAT5, as indicated by the motif analyses. Although higher-resolution mapping of *PAN3-AS1*-associated chromatin domains and nuclear contacts would help refine this mechanistic model, such analyses remain technically challenging (Goldrich et al., 2026), precluding a definitive understanding of this aspect of the mechanism within the current study.

A major translational barrier for RNA-based therapeutics is the efficient and selective delivery of therapeutic payloads to disease-relevant tissues. This challenge is particularly significant in leukemia, where malignant cells are distributed across the circulation, bone marrow, spleen, and other hematopoietic compartments. Most clinically advanced lipid nanoparticles preferentially accumulate in the liver, limiting their application to extrahepatic malignancies. Several studies have reported efforts to develop and apply LNP technology for leukemia; however; most have been limited to *in vitro* or *ex vivo* testing mainly with cell line models and demonstrated limited specificity and efficacy (Hiraki et al., 2026; Jyotsana et al., 2019; Sørensen et al., 2025). Here, we reported an LNP^HiPEH^ formulation enabling efficient delivery of siRNAs into AML cells and produced sustained suppression of the target *PAN3-AS1* lncRNA *in vivo*, resulting in substantial disease control in experimental animals. These superior results open opportunities to evaluate other coding and non-coding RNA dependencies that have previously been considered difficult to drug, particularly in hematological malignancies. Using *PAN3-AS1* as a case study, we demonstrate that lncRNAs may represent an attractive and targetable therapeutic class with a very favorable therapeutic window and limited toxicity in normal hematopoietic cells and non-hematopoietic tissues. Future toxicological and pharmacological studies will further define the safe and optimal dosing schedule for improved efficacy.

In addition, with the identified functional link between *PAN3-AS1* and FLT3 activation, we propose a combination treatment of LNP-siPAN3-AS1 and FLT3 inhibitors. FLT3 mutations are among the most frequent genetic alterations in AML and are associated with aggressive disease and a high risk of relapse. The enhanced and sustained antileukemic effects observed when *PAN3-AS1* inhibition was combined with FLT3-targeted therapy imply that disruption of the *PAN3-AS1–FLT3* regulatory module may increase the depth or durability of response. This strategy may complement direct kinase inhibition by simultaneously reducing FLT3 expression and attenuating additional leukemia-supporting programs controlled by *PAN3-AS1*. Moreover, although the strongest rationale for combination therapy is currently in FLT3-mutant AML, *PAN3-AS1* therapeutic intervention is not restricted to this molecular subtype. The elevated expression of *PAN3-AS1* across multiple AML genetic backgrounds—including NPM1-mutant, IDH1/2-mutant, and KMT2A-rearranged leukemias— suggests that this lncRNA may represent a shared regulatory node across different oncogenic contexts. This convergence is consistent with the observation that *FLT3* is itself a transcriptional target of the HOXA9/MEIS1 axis, a core leukemogenic program downstream of both NPM1 mutations and KMT2A rearrangements (Zhou and Lu, 2023). Its elevated expression across additional AML groups raises the possibility that *PAN3-AS1* inhibition could have broader therapeutic activity, either as a single agent or in combination with subtype-specific therapies. Future studies will further define the genetic and molecular features that predict sensitivity to *PAN3-AS1* depletion. Such analyses may also nominate rational combinations with chemotherapy, epigenetic therapies, or other targeted agents.

Importantly, our findings add to growing evidence that long non-coding RNAs embedded within oncogenic loci can act as regulatory elements that propagate leukemic signaling beyond their site of origin (Cheng et al., 2021; Cho et al., 2018; Gupta et al., 2010; Lyu et al., 2026; Papaioannou et al., 2019; Trimarchi et al., 2014; Yildirim et al., 2013). More broadly, we illustrate how non-coding transcription can cooperate with recurrent genetic alterations to reinforce oncogenic signalling. The genomic proximity of *PAN3-AS1* to FLT3 suggests that genetic and non-coding regulatory mechanisms may form a feed-forward circuit in which aberrant signalling promotes a leukemia-associated transcriptional state that, in turn, sustains expression or activity of the mutated oncogene. Similar relationships may exist at other cancer-associated loci, as many lncRNAs are located near genes that are recurrently mutated, amplified, translocated, or otherwise dysregulated in malignancy. Non-coding transcripts arising from these regions should therefore not be assumed to represent transcriptional noise. Instead, they may function as integral components of oncogenic regulatory circuits and create therapeutic vulnerabilities that are distinct from those associated with the neighbouring protein-coding gene.

In conclusion, this study identifies *PAN3-AS1* as a critical regulator of AML survival and establishes a functional link between a leukemia-associated lncRNA, chromatin organization, and oncogenic signalling. Our preclinical data demonstrating that *PAN3-AS1* can be therapeutically targeted using systemically delivered siRNA-loaded LNPs provides a strong proof of principle for translating lncRNA biology into RNA-based treatment strategies. The work expands the biological understanding of non-coding regulatory circuits in cancer and supports the potentially generalized interest in investigating lncRNAs located near recurrent cancer-associated genetic alterations as both mechanistic mediators and therapeutically exploitable targets.

## Acknowledgments

The work was supported by the Terry Fox Research Institute New Investigator Award to L.P.V., and Michael Smith Health Research BC Trainee Award and BC Cancer Rising Star to Z.J. L.P.V is the Scholar of the American Society of Hematology, Scholar of the V foundation for Cancer Research and is a Tier 2 Canada Research in RNA biology and Hematological Malignancies and is supported by the Michael Smith Health Research Scholar award. M.S. is an FRQ Research Scholar and hold the Low-Beer Family Research Chair on Noncoding RNA Biology & Therapeutics. L.P.V’s lab is supported by Canadian Institutes for Health Research Project Grant (CIHR), Natural Sciences and Engineering Research Council of Canada Discovery Grant and the Terry Fox New Frontiers Program Project Grant - 1135 (TFRI PPG 1135). M.G.K. is a Scholar of the Blood Cancer United (Formerly Leukemia Lymphoma Society and Basic Science Discovery Grant, and was supported by NIDDK NIH R01-DK101989-01A1, NCI 1R01CA193842-01, R01HL135564, R01CA274249-01A1, R01CA186702, R01CA283578, and R01CA225231-01. D.L, D.G, and J.G are supported by TFRI PPG 1135 and CIHR PJT-162131. The LNP development and formulation work performed at P.R.C’s lab was funded by the Canada First Research Excellence Fund awarded to the D2R Initiative at McGill University, the Canada Biomedical Research Fund, the Biosciences Research Infrastructure Fund, and Genome British Columbia (B31AVG). We thank the staff of Life Science Imaging Core in University of British Columbia; Canada’s Michael Smith Genome Sciences Center (GSC), Flow Core, Animal Research Center at BC Cancer for their technical assistance. We thank the flow cytometry, transgenic core facility at Memorial Sloan Kettering Institute, which are supported by the core Grant P30 CA008748. We are grateful to our lab members, Dr. Connie Eaves lab members, Dr. Junbum Im, Dr. Martin Hirst, Dr. Andrew Wong for their discussion and technical assistance.

## Author contributions

Conceptualization, Z.J., M.G.K., and L.P.V.; methodology, Z.J., K.Y.T.C., S.H., G.B.M., A.K., G.B., P.R.C., and L.P.V.; investigation, Z.J., B.M., A.L., J.L.C., Y.L., S-W.G.C., S.E.S., P. E.F.S., G.E., F.W., K.D., Q.A.H., T.B., A.S., K.D., T.C., F.P., and L.P.V.; formal analysis, Z.J., A.L., L.S., D.L., J.G., D.G., Z.L., L.E., O.C., M.S.; resources, F.K., M.G.K., A.K., P.R.C., and L.P.V.; writing – original draft, Z.J., and L.P.V.; writing – review & editing, Z.J., K.Y.T.C, M.S, F.P., M.G.K, and L.P.V.; supervision, L.P.V.; funding acquisition, M.G.K, A.K, P.R.C and L.P.V.

## Resource availability

### Lead contact

Further information and requests for resources and reagents should be directed to and will be fulfilled by the lead contact, Ly P. Vu.

### Materials availability

Cell lines newly generated in this work are available upon request to the lead contact.

### Data and code availability

No new code is generated through this study. The complete sequencing datasets are available at NCBI PRJNA1495346.The reviewer link is: https://dataview.ncbi.nlm.nih.gov/object/PRJNA1495346?reviewer=6g7s9ojufmuiovoate8ts351ju.

## STAR★Methods

### Experimental model and study participant details

#### Human primary cord blood and patient samples

Human cord blood (CB) samples were obtained from Eaves Stem Cell Assay Laboratory in the Terry Fox Laboratory in the BC Cancer Agency or purchased from STEMCELL Technologies. Human primary AML patient samples were obtained from HEMATOLOGY CELL BANK of BC. Cryopreserved samples were thawed in media containing 4.5 mL Iscove’s Modified Dulbecco’s Medium (IMDM, Thermo Fisher Scientific, #12440061), 4.5 mL Fetal Bovine Serum (FBS, Thermo Fisher Scientific, # 12483020) and 1% DNase I (Sigma Aldrich, #D4513-1VL, 10 mg/mL). Cells were cultured in serum-free media (80% IMDM, 20% BIT 9500 (STEMCELL Technologies, #09500, add 0.2mg LDL for 1mL BIT), 100 nM 2-Mercaptoethanol (Sigma Aldrich, # M6250-100ML) and 2 mM L-Glutamine (Thermo Fisher Scientific, # 35050061)) supplemented with 5 growth factors (20 ng/mL hIL-3 (Thermo Fisher Scientific, #200-03-50UG), 20 ng/mL hIL-6 (STEMCELL Technologies, #78148), 100 ng/mL hSCF (STEMCELL Technologies, #78062), 20 ng/mL G-CSF (STEMCELL Technologies, #78012), 100 ng/mL rhFLT3-ligand (STEMCELL Technologies, #78009). To preserve stemness of CD34+ CB cells, 35 nM UM171 (STEMCELL Technologies, #72912) and 750 nM StemRegenin 1 (STEMCELL Technologies, #72354) were added to the 5-growth factor serum-free media. The cells were incubated at 37 °C in humidified 5% CO_2_ incubator. All studies were performed according to human ethics H24-02079 approved by the Research Ethic Board at the University of British Columbia.

#### Human Cell lines

Parental human leukemia cell lines MOLM13, MV4-11, OCI-AML3, THP1, KG-1, KCL-22, Kasumi, K562, and HL-60, as well as transgene-expressing cells generated in this study, were cultured in RPMI 1640 medium (Thermo Fisher Scientific, #11875119) containing 10% FBS. HEK293T and HeLa cells were cultured in Dulbecco’s Modified Eagle Medium (DMEM, Thermo Fisher Scientific, #11875119) medium containing 10% FBS. All the cells were incubated at 37°C with 5% CO2 incubator and routinely tested for mycoplasma contamination. All human cell lines were previously purchased from ATCC and were authenticated via STR profiling using Genetica cell line testing service.

#### Mouse cells

RN2 mouse leukemia cell line cells were originally obtained from ATCC and cultured in RPMI 1640 medium containing 10% FBS. Primary mouse bone marrow cells, and mouse-derived MLL-AF9 leukemia cells were cultured in StemSpan™ Serum-Free Expansion Medium (SFEM, STEMCELL Technologies, #09650) supplied with mouse cytokines, 10ng/mL rmIL-3 (STEMCELL Technologies, #78042.1),10ng/mL rmIL-6 (STEMCELL Technologies, #78052.1), 50ng/mL rmSCF (CedarLane, #250-03-100UG), 10ng/mL rmTPO (CedarLane, #315-14-10UG) and 20ng/mL rmFlt3l (CedarLane, #250-31L-50UG). Cells were incubated at 37°C with 5% CO_2_ incubator.

#### Mice

C57BL/6 and Pep3b female mice (expressing CD45.1, congenic to C57BL/6) (expressing CD45.2) used for transplant assays were purchased from BCCRC Animal Research Centre. Lnc35682 homozygous knock out mice were generated in C57BL/6 in house. Both male and female mice were used for experiments with age- and gender-matched littermates as control. NSG (NOD-scid IL2Rgnull), NRG-3GS (NOD.Rag1−/−;γcnull-IL3/GM/SF) used for xenograft assays were purchased from BCCRC Animal Research Centre. Female mice were used for experiments with age-matched littermates as control. All studies were performed according to animal protocol # A23-0148 approved by the Animal Care and Use Committee at the University of British Columbia.

### Method details

#### Plasmids

For shRNA-mediated depletion, SGEP-puromycin-EGFP miR-E shRNA constructs were used to target PAN3-AS1. shRNA sequences are listed in Table S1. The plasmid DNA was used as the transfer vector for lentiviral production.

For CRISPR/CasRx mediated RNA silencing, EF1a-CasRx-2A-EGFP (Addgene#109049) were used to generate CasRx expressing cell lines. sgRNAs for PAN3-AS1 targeting were designed using Cas13design tool (Guo et al., 2021; Wessels *et al*., 2020). Oligos were synthesized from IDT and cloned into CasRx-guideRNA-puromycin backbone (Addgene#134839). Guide RNA sequences are listed in **Table S1**. The plasmid DNA was used as the transfer vector for lentiviral production.

For *Lnc35682* overexpression, the *de novo* assembled *Lnc35682* sequence was synthesized and cloned into the pMSCV-EBFP vector at Azenta Life Sciences, (formerly Genewiz), with the empty backbone used as the control. The plasmid DNA was used as the transfer vector for retroviral production. For PAN3-AS1, FLT3-ITD, and PAN3 overexpression, sequences were synthesized and cloned into the pMNDU3-pgk-EYFP vector at Azenta Life Sciences, with the empty backbone used as the control. Truncated *PAN3-AS1* constructs were subcloned into the same vector. The plasmid DNA was used as the transfer vector for lentiviral production.

For creating luciferase-expressing cells, MPAP-PGK-CBR-YFP and MPAP-PGK-CBG99-GFP constructs were used for making lentiviruses. The constructs were gifts from Dr. Connie Eaves lab. The plasmid DNA was used as the transfer vector for lentiviral production.

For lentiviral and retroviral production, pMD2.G, pCMVd8-78-delta R, pRSV-REV, psPAX, PCL-ECO, and PCL-AMPHO were used as envelope and packaging plasmids.

#### CRISPR/Cas9 Knockout mouse

The CRISPR gRNAs used to delete *Lnc35682* were designed using the approach of Romanienko et al (Romanienko et al., 2016; Vu et al., 2017). Both were produced by in vitro transcription using the pU6T7 promoter in the hybrid plasmid described. To initiate cleavage of the target locus in mice, gRNAs in conjunction with Cas9 mRNA were co-injected into the pronuclei of mouse zygotes at a concentration of 50ng/μl each, using conventional techniques. Deleted samples were assayed using PCR primers Lnc35682-F1,F4, and R1 listed in Table S1. The size of the deletion based on nucleotide length of the amplicon obtained ∼260bp for wild type allele versus ∼ 500bp for the *Lnc35682* knock out allele.

#### Virus production

For lentiviruses production, H293T cells were seeded in 10 cm plates and grown to 70-80% confluency before cell transfection. The lentiviral transfer vector plasmid, packaging plasmid (psPAX2), and envelop plasmid (pMD2.G) were combined at 4:3:1 ratio (12.5 μg:9.375 μg:3.125 μg, respectively) in 1 mL of 0.25 M CaCl_2_. Alternatively, using lentiviral transfer vector plasmid, pCMVd8-78-delta R, pRSV-REV, and VSV.G (10µg:6.5µg:2.5µg:3.5µg). For retroviruses production, to transfect mouse cells, transfer plasmid, packaging plasmid (PCL-ECO), and envelop plasmid (pMD2.G) were combined at 4:3:1 ratio (12.5 μg:9.375 μg:3.125 μg, respectively) in 1 mL of 0.25 M CaCl_2_. To transfect human cells, transfer plasmid, packaging plasmid (PCL-AMPHO), and envelop plasmid (pMD2.G) were combined at 4:3:1 ratio (12.5 μg:9.375 μg:3.125 μg, respectively) in 1 mL of 0.25 M CaCl_2_. Add the mixture to 1 mL of filtered 2× concentrate BES buffered saline (BioWORLD, #40220006-2) in a dropwise manner. A sterile 1 mL pipette was used to gently create bubble air through the DNA mix. Following the incubation at room temperature for 5–10 mins. the solution was added dropwise to H293 T cells. The plates were rocked gently in a circular motion to distribute the precipitates. Cells were cultured at 37 °C in a humidified 5% CO_2_ incubator for 16 hours, followed by replacement with fresh media. Viral supernatants were collected twice at 24 hours and 48 hours after replenishing new media. The supernatant was passed through a 0.45 μm pore PVDF Millex-HV filter (Millipore, SLHVR33RB) and stored at 4°C for less than 2 weeks or −80 °C. Ultracentrifugation can be optionally performed to concentrate viral particles from the supernatant by approximately 100-fold.

#### Viral transduction and cell selection

Leukemia cell lines were plated in RPMI with 10% FBS at a density of 5 × 10^5^ cells/mL with 5μg/mL polybrene. Cells were transduced with lentiviruses or retroviruses by spinfection at 1400 × g, room temperature for 1 hour. Replace the medium 16h after transduction. Cells were selected 48 h after transduction by adding 3 µg/mL puromycin or by FACS sorting of fluorophore-positive cells.

CD34+ cord blood cells were thawed and seeded at a density of 1 x 10^6^/mL for 16h with 8μM Cyclosporin H (Toronto Research Chemicals, # C988920-2) in 96-well round bottom plates. Cells were then incubated with concentrated lentiviruses (1μl per well, titers ∼10^9^/mL) for 6 h, after which the medium was replaced with fresh medium. Cells were selected 48 h after transduction by adding 3 µg/mL puromycin or by FACS sorting of fluorophore-positive cells. Collected cells were then subjected to downstream assays, including liquid culture, colony-forming assay, and *in vivo* transplantation.

Primary AML cells, either thawed or freshly isolated PDX cells from leukemic mice, were seeded at a density of 1 × 10⁶ cells/mL in 96-well round bottom plates. Cells were then incubated with concentrated lentiviruses (1μl per well, titers ∼10^9^/mL) for 16 h, after which the medium was replaced with fresh medium. Cells were selected 48 h after transduction by FACS sorting of fluorophore-positive cells. Collected cells were then subjected to downstream assays, including liquid culture, colony-forming assay, and *in vivo* transplantation.

To generate MF9-leukemia cells, C57BL/6 wild-type (WT) or Lnc35682 CRISPR-KO (KO) mouse bone marrow cells were isolated and subsequently enriched for c-Kit+ cells with magnetic beads separation using AutoMACS. c-Kit+-enriched cells were stained with a lineage antibody cocktail (antibodies to CD3, CD4, CD8, Gr-1, B220, CD19, and Ter119 conjugated with PeCy5) and with Sca–Pac Blue, CD34-FITC, SLAM-APC, CD48-PE, and c-Kit-APC-Cy7 antibodies. LSK cells were sorted using a BD FACSAria instrument. Sorted cells were grown overnight in retronectin coated round bottom 96-well plate. Cells were transduced twice with supernatant containing retroviruses encoding MLL-AF9 and GFP (a gift from S. Armstrong, Memorial Sloan Kettering Cancer Center) together with 4μg/mL polybrene for 24 hours. The cells were expanded for 1 week in MethoCult GF M3434 (StemCell Technologies, #03434). Cells were sorted for GFP positivity for subsequent experimental usage.

#### Colony Forming Assay

CFC assays for human cells were performed in a semi-solid methylcellulose (MethoCult, StemCell Technologies, #04434). 500 −1,000 CB-CD34+ cells or 30,000 PDX cells were plated in six-well plate (in duplicate) containing 1 mL methylcellulose. Plates were kept in incubator at 37 °C and 5% CO_2_. Erythroid progenitor cells (E), granulocyte-macrophage progenitor cells (GM), and multipotent granulocyte, erythroid, macrophage, and megakaryocyte progenitor cells (GEMM) colonies were scored under a microscope after 14 days. For mouse cells, 10,000 cells were plated on MethoCult GFM3434. Colonies were scored every 5 days for leukemia cells and every 7 days for normal c-Kit-enriched bone marrow cells.

#### RNA extraction and quantitative real-time PCR assays (qRT-PCR)

Total RNA was extracted from cells using RNA Purification Kit (NORGEN, #48702) following the standard manual. An equal amount of RNA from samples were reverse transcribed into cDNA with iScript™ Reverse Transcription Supermix (Biorad, 1708841), and qPCR was performed using a QuantStudio™ 5 Real-Time PCR System detection using primers together with PowerTrack™ SYBR Green Master Mix (Thermo Fisher Scientific, #A46111). Reactions were run in a 10 μl reaction according to the manufacturer’s protocol in triplicates. Cycling conditions included an initial hold step (95 °C for 10 seconds) and 40 cycles of a two-step PCR (95 °C for 15 seconds and then 60 °C for 60 seconds), followed by a dissociation step (95 °C for 15 seconds). Relative messenger RNA (mRNA) expression was calculated by the comparative 2^−ΔΔCT^ method. GAPDH or ACTIN was used as an internal control.

#### Immunoblot analysis

For immunoblot analysis, cells were counted and washed twice with cold PBS prior to collection. Cells were then lysed in 1× Laemmli protein loading buffer and boiled at 95 °C for 5 minutes. Cell lysates were run on a 4%–15% Mini TGX precast protein gels or 4–15% Criterion™ TGX™ Precast Midi Protein Gel (Biorad, #4568084, 4568085, 4568086, 5678084) and transferred to a nitrocellulose membrane (Biorad, #1620115). Membranes were blocked with 5% milk for 1 hour prior to incubation with antibodies against ACTIN (Sigma Aldrich, A3854, 1:5000), FLT3 (NEB, #3462S), p-FLT3 (NEB, #60413S), STAT5 (NEB, #25656T), p-STAT5 (NEB, #9314S), LMNB1 (Abcam, #ab16048), HA (NEB, # 3724S). After overnight primary antibody incubation, the membranes were washed with 1 × PBST and then incubated with HRP-linked secondary antibodies (goat anti-mouse IgG [NEB, 7076 V] or goat anti-rabbit IgG [NEB, 7074 V], 1:3000) for 1 hour at room temperature. Immobilon ECL Western HRP Substrate (Millipore, WBKLS0500) and ECL Western Blotting Detection Reagent (Sigma, GERPN2105) were used to detect the protein bands on the Bio-Rad ChemiDoc Imaging System with chemiluminescence detection. The relative protein expression of each protein band was analyzed by Image J software, using ACTIN as the reference.

#### Immunofluorescence and RNA-Fluorescence In Situ Hybridization (FISH)

Cells were fixed in 10% neutral buffered formalin (VWR, #77556) at a concentration of 1x 10^6^ cells/mL for 60 minutes at 37 °C. They are then resuspended in 70% Ethanol. Cytospin 1 × 10^5^ cells onto a slide at 750 × g for 5 minutes using the ThermoFisher Cytospin 4. Cells are then permeabilized with 50%, 70%, and 2 × 100% Ethanol for 5 minutes each. After outlining the cell spot with a hydrophobic pen, cells are incubated in 0.5% PBST blocking solution for 60 minutes at room temperature. For FISH, cells were then incubated with RNA-scope probes (ACD bio-techne, #1190621-C3) for 120 minutes at 40 °C. Consequent FISH signal amplification steps were performed according to the protocol outlined in the RNAScope Assay kit from ACD bio-techne. Next, cells were incubated in primary antibody anti-LMNB1 (Abcam, #ab16048) 1:500 in the blocking solution (Abnova, #H00004849-M01) overnight at 4 °C. Cells were then washed and incubated with secondary antibody anti-rabbit AF647: 1:500 (A32795) for 60 minutes at room temperature. Finally, cells were washed again and mounted in mounting medium with DAPI (SIGMA, #DUO82040). Slides were stored at −20 °C for 24 hours before imaging on the Leica SP8 gSTED Confocal Microscope. Images were analyzed using LAS X Office and organized in Adobe Illustrator.

#### Flow cytometry and Fluorescence-activated Cell Sorting (FACS)

To assess engraftment of human leukemia cells in NSG/NRG-3GS recipient mice, blood and bone marrow cells collected from animals were treated with red blood cell (RBC) lysis buffer (NEB, #46232 S) prior to staining with two antibodies against human CD45 surface marker i.e., anti-hCD45-AF700 clone 2D1 (Biolegend, #368514) and anti-hCD45-PB clone H130 (Biolegend, #304029). To analyze cell lineage, CD34, CD38, CD11b, CD10, CD19, CD3, CD71, CD135 were used as different lineage markers. Refer to Key Resource Table.

To analyze mouse cell lineage, lineage antibody cocktail (Lin-: antibodies to CD3, CD4, CD8, Gr-1, B220, CD19, and Ter119 conjugated with PeCy5) and with Sca1–Pac Blue, c-Kit-APC-Cy7, CD34-FITC, CD150-APC, and CD48-PE antibodies were used for analysing mouse normal LT-HSCs (CD150+CD48-LSK), GMPs (CD34+CD32/16mid Sca-1-c-Kit+) and LSCs (c-Kit^high^). Flow analysis on CD45.1 and CD45.2 surface markers from cells isolated from doner mice bone marrow were used for *in vivo* chimerism ratio to determine repopulation ability of the donor cells.

DAPI (SIGMA, #D8417-5MG), Propidium Iodide, (Thermo Fisher Scientific, #P1304MP), or Zombie Viability Kits (Biolegend, #423125, 423101) were used for live/dead cell gating. Flow cytometry samples were analyzed on a BD FACS LSR Fortessa, Symphony A1, Symphony A5, or Symphony A5 SE instruments. FACS were done on sorters BD Aria Fusion or BD Aria III.

#### *In vivo* transplantation and functional readout

2 x 10^5^ MF9 leukemia cells carrying scrambled shRNA or shRNA targeting *Lnc35682* were injected retro-orbitally with 2.5 x 10^5^ helper bone marrow cells into lethally irradiated 6-8-week-old C57BL/6 mouse. 2 x 10^5^ MF9 leukemia cells derived from wild type or Lnc35682 knockout mice and 2.5 x 10^5^ helper bone marrow cells were injected retro-orbitally into each lethally irradiated 6- to 8-week-old C57BL/6 mouse to assess disease progression ability.

Pep3b female mice expressing CD45.1, congenic to C57BL/6, were used as recipients for transplant assays. Recipients were lethally irradiated with 2 doses of 5 Gy using a cesium irradiator and injected with 2 x 10^6^ total bone marrow cells isolated from the WT or Lnc35682 knockout donor mice (C57BL/6, expressing CD45.2). Peripheral blood and bone marrow were then monitored over 24 weeks for CD45.2 chimerism to determine repopulation ability of the donor cells via flow cytometry.

#### *In vivo* xenograft of AML cells and patient-derived xenografts (PDXs)

For cell line cells, 3 x 10^5^ luciferase-expressing MOLM13 cells carrying either scrambled shRNA or shRNA for *PAN3-AS1* knockdown were tail vein injected to NSG mice. 2 x 10^5^ luciferase-expressing MOLM13 cells carrying either empty vector or *PAN3-AS1* OV cells were tail vein injected to NSG mice. Female NSG mice were used at age 9-12 weeks old with sub-lethal irradiation (350cGy) before transplantation. 8 x 10^4^ - 1 x 10^5^ luciferase-expressing MV4-11 cells were tail vein injected to NRG-3GS mice before LNP or drug administration.

For primary cells, 5 x 10^4^ CD34+ cord blood cells carrying either scrambled shRNA or shRNA for *PAN3-AS1* knockdown were tail vein injected to NRG-3GS mice. For AML patient cells, variable number of cells were tail vein injected to develop patient derived xenograft (PDX) models. After secondary or tertiary PDX expansion, 3 × 10^5^ PDX cells carrying either scrambled shRNA or shRNA for *PAN3-AS1* knockdown were tail vein injected to NRG-3GS mice; 3 × 10^5^ – 1x 10^6^ luciferase expressing PDX cells were tail-vein injected into NRG-3GS mice before LNP and drug administration. Female NRG-3GS mice were used at age 9-12 weeks old with lethal irradiation (800cGy) before xenograft.

For *in vivo* therapeutic treatment models, LNP-siRNA (2.5 mg/kg) was administered to recipient mice by tail vein injection twice per week. Gilteritinib (10 mg/kg) was administered by oral gavage three-times per week, on alternating days. Treatments were initiated between 24 hours to 1 week post-transplantation and continued until the mice reached the experimental endpoint. For PDX-FLT3 mutant AML model, treatments were initiated 4 weeks post-transplantation. Mice were purchased from BCCRC Animal Research Centre, and procedures all complied with Animal protocols approved by the University of British Columbia.

#### In Vivo Imaging System (IVIS)

D-luciferin (GoldBio, #LUCK-1G) was freshly dissolved in PBS at 15 mg/mL and sterile-filtered through a Cytiva Whatman™ Vacu-Guard filter (Thermo Fisher Scientific, #09-744-76). Mice were intraperitoneal injected with 200 μL per mouse, and bioluminescence images was captured 11 min after injection on IVIS Lumina S5 machine following manufacture’s instruction at auto-exposure or fixed time exposure. Data was quantified and analyzed on Living Image 4.7.4 software.

#### Transcriptomic profiling (RNA-seq)

Total RNA isolated from biological samples was processed with Poly(A) selection and library preparation using TruSeq RNA Library Prep Kit v2 (Illumina, RS-122-2001). Each group has three biological repeats. Libraries were sequenced with 2 × 150 bp paired-end run and, in total 20–40 million reads/sample using Illumina® NovaSeq.

#### RNA-seq data analysis

Raw sequencing reads were subjected to quality control and adapter trimming using FASTP (v0.23.4). Clean reads were then aligned to the reference genome (GRCh38/hg38) using STAR (2.7.10b). Gene-level read counts were generated using featureCounts (v2.0.3) based on the corresponding gene annotation file. The resulting count matrix was imported into R for downstream analysis. Differential gene expression analysis between experimental groups was performed using DESeq2. Genes with an adjusted p-value < 0.05 and log_2_fold change ≥ 1 or < −1, were considered significantly differentially expressed. Normalized count data were used for downstream visualization and statistical analyses.

#### Assay for transposase-accessible chromatin using sequencing (ATACseq)

50,000 live cells from each biological sample were used for ATAC-seq library preparation using the ATAC-seq Kit (Active Motif, #53150) according to the manufacturer’s instructions. Briefly, cells were lysed to isolate nuclei, followed by Tn5 transposase-mediated tagmentation to simultaneously fragment genomic DNA and insert sequencing adapters at accessible chromatin regions. The transposed DNA fragments were then PCR-amplified to generate sequencing libraries, followed by library purification and size selection. Library quality and concentration were assessed prior to next-generation sequencing. Each group has three biological repeats. Libraries were sequenced with 2 × 150 bp paired-end run and, in total 20–40 million reads/sample using Illumina® NovaSeq.

#### ATACseq data analysis

The paired-end reads were processed with the ENCODE ATAC-seq pipeline v2.2.3 using default parameters. As a part of the pipeline, reads were aligned to the hg38 reference genome using Bowtie2 v2.3.4.3 and consensus peaks were called using MACS2 v2.2.4(Langmead *et al*., 2019; Zhang *et al*., 2008). Differential peak calling was performed with HOMER v4.11 using DESeq2 v1.34 with three replicates for each condition(Love *et al*., 2014). A peak is considered differentially accessible if it has a log_2_fold change magnitude > 1 and a Benjamini-Hochberg adjusted p-value > 0.05. The differential peaks were then analyzed for known and *de novo* motifs using HOMER (*findMotifsGenome.pl -size 200*), and annotated with hg38 transcription factor binding sites from the LOLA database v170206 using annotatr v1.20(Cavalcante and Sartor, 2017; Sheffield and Bock, 2016).

#### RNA immunoprecipitation qPCR (RIP-qPCR)

1 × 10^7^ cells were used for RNA immunoprecipitation optimized from a previously described protocol(Baker et al., 2023). Cells were washed with cold PBS and then lysed in 100μl lysis buffer (50mM Tris-HCL, pH 7.5, 1mM EDTA, 150mM NaCl, 0.5% NP-40), with freshly added 100x Proteinase Inhibitor (Thermo Fisher Scientific, #78429), 5U/mL RNase Inhibitor Superase In (Thermo Fisher Scientific, #AM2696), 5μl/mL 0.1M PMSF. 4.5μg of anti-LMNB1 antibody (Abcam, #ab16048) or anti-rabbit IgG (Thermo Fisher Scientific, #10500C) incubated overnight with 20μl magnetic beads (Thermo Fisher Scientific, #10003D) to immunoprecipitated LMNB1 or non-specific bindings. After the immunoprecipitated complexes were washed with lysis buffer. RNA extraction was performed by the phenol–chloroform method, and purified RNA was converted to cDNA using iScript™ Reverse Transcription Supermix (Biorad, 1708841), and qPCR was performed to validate reciprocal interactions.

#### siRNA Transfection

Negative control siRNA, positive control siRNA and transfection control labeled with TYE563 were obtained by ordering DsiRNAs in a TriFECTa® RNAi Kit (IDT). Custom siRNAs targeting *PAN3-AS1* or CBR/CBG99-luciferase was synthesized at IDT, see **Table S1**. Leukemia cells were plated in a density of 2×10^5^/mL into plates coated with Geltrex (Thermo Fisher Scientific, # A1413301) and incubate overnight before transfection. For Geltrex coating, 10μL Geltrex was diluted in DMEM, and 400μL coating solution was added to each well of a 24-well standard plate, incubating overnight at 37°C. siRNA was transfected using Lipofectamine™ RNAiMAX Transfection Reagent (Thermo Fisher Scientific, #13778030) according to the manufactural instructions. For LNP encapsulated siRNA transfection, LNP-siRNAs were diluted in PBS and added to the cell culture prepared similarly to the Lipofectamine transfection. Cells at 24h, 48h, or 72h post transfection were collected for transfection efficiency, gene silencing efficacy, and cell proliferation assessments.

#### Drug treatment

Quizartinib (AC220, Selleckchem, #S1526) or Gilteritinib (Selleckchem, #S7754) was dissolved in DMSO (Sigma, #D2650-100ML) and subjected to 10-fold serial dilutions for in vitro treatment. Cell proliferation was assessed 72 h after treatment using either a cell counter or the CellTiter-Glo® 2.0 assay (Promega, #G9242), according to the manufacturer’s instructions.

For *in vivo* treatment, 2 mg Gilteritinib was dissolved sequentially in 3% (v/v) anhydrous ethanol and 5% (v/v) DMSO, followed by 20% (v/v) PEG300, 5% (v/v) Tween-80, and 67% (v/v) PBS, to a final volume of 1 mL, in that order. The solution was mixed thoroughly at each step to ensure complete dissolution and the absence of precipitation. The vehicle solution was prepared using the same formulation without Gilteritinib.

#### Synergy assay

Combination treatment of Gilteritinib with LNP formulated siRNA targeting *PAN3-AS1* were performed to evaluate the synergistic effects of FLT3 inhibition and *PAN3-AS1* silencing. Gilteritinib was treated with doses 0, 0.3, 0.6, 1, 3, 6nM. LNP formulated siRNA was treated with doses, 0, 0.5, 1, 3, 5μg/mL. 3 replicates were done for each condition. Cell proliferation was assessed 72 h after treatment using CellTiter-Glo® 2.0 assay (Promega, #G9242), according to the manufacturer’s instructions. Data was analyzed in SynergyFinder (https://synergyfinder.org).

#### Luciferase Assay

Cells were lysed in RIPA buffer (Thermo Fisher Scientific, #89901) on ice according to the manufacturer’s instructions. Cell lysates were divided into two portions: one for protein quantification using the Pierce™ BCA Protein Assay Kit (Thermo Fisher Scientific, #23227) and the other for the luciferase assay using the Bright-Glo™ Luciferase Assay System (Promega, #E2610) according to the manufacturer’s instructions. Signals were captured on a TECAN plate reader. Luciferase activity was normalized to protein levels based on the corresponding BCA assay results.

#### Myeloid leukemia targeting LNP formulation

##### Materials

The ionizable lipid, (9Z,12Z)-1-(8Z,11Z)-8,11-heptade butanoic acid, 4-(dimethylamino)- cadien-1-yl-9,12-octadecadien-1-yl ester (nor-MC3), was synthesized by the laboratory of Dr. Glenn Sammis at the University of British Columbia. A mix of sphingomyelin lipids extracted from chicken egg (ESM), and 1,2-dimyristoyl-rac-glycero-3-methoxypolyethylene glycol-2000 (PEG2k-DMG), were purchased from Avanti Research (Alabaster, AL, USA). Cholesterol and Dulbecco’s Phosphate Buffered Saline (DPBS) were purchased from MilliporeSigma (Burlington, MA, USA). Citric acid monohydrate was purchased from BioShop Canada Inc. (Burlington, ON, Canada). Sodium hydroxide was purchased from Thermo Fisher Scientific (Waltham, MA, USA).

##### Formulation of Lipid Nanoparticles

Lipid nanoparticles (LNPs) were formulated as per previously reported methods (Jeffs et al., 2005), and the composition was taken from a study on the extrahepatic delivery of messenger RNA via an LNP platform (*manuscript under review*). In brief, norMC3, ESM, cholesterol, and PEG2k-DMG were dissolved in ethanol. The total lipid concentration was 10 mM. Sodium citrate buffer was made by dissolving citric acid monohydrate in water at a concentration of 300 mM, and then adjusting the pH to 4.0 by the dropwise addition of sodium hydroxide solution. This buffer was further diluted and mixed with the small interfering RNA (siRNA). The organic phase containing the lipids was then mixed with the aqueous phase containing the siRNA using a T-junction microfluidics mixer at a volume ratio of 1:3 and a total flow rate of 20 mL/min. The charge ratio between the cationic groups in the ionizable lipid and the anionic groups in the siRNA (N:P ratio) was 3:1. After mixing the two phases, the LNP suspension was dialyzed overnight at room temperature in DPBS, pH 7.4. It was then passed through a 0.2 µm syringe filter (Cytiva Life Sciences) and concentrated down to an encapsulated siRNA concentration of 0.5 mg/mL using 10 kDa NMWL Amicon Ultracentrifugal filter units (MilliporeSigma). The LNPs were stored at 4°C for no longer than 2 weeks until use. Characterization of lipid nanoparticles: Quantification of siRNA was done using a Quant-it RiboGreen Assay (Thermo Fisher Scientific) according to the manufacturer’s instructions. LNPs were divided into two fractions, one left untreated, and another treated with Triton X-100 (MilliporeSigma) at 2% volume per volume. LNPs treated with Triton X-100 would release the siRNA payload. The RiboGreen reagent was then added to the LNPs and the free siRNA standards. All the samples were read on a Spark multimode microplate reader (Tecan Group Ltd., Männedorf, Switzerland) at an excitation wavelength of 500 nm and an emission wavelength of 525 nm. The siRNA concentration in the LNP samples was calculated by comparing the fluorescence signal to that obtained by the free siRNA standards. Encapsulation efficiency was calculated by the following formula:

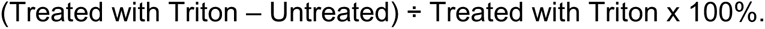

Cholesterol content was measured using a Wako Cholesterol E assay kit (Fujifilm Healthcare Solutions, Lexington, MA, USA) according to the manufacturer’s instructions. Cholesterol standards and the LNP samples were read on a Spark multimode microplate reader (Tecan Group Ltd., Männedorf, Switzerland) at an absorbance wavelength of 595 nm.

LNP hydrodynamic diameter and polydispersity index were measured using a Nanosizer Lab dynamic light scattering analyzer (Malvern Panalytical, Malvern, Worcestershire, UK) according to manufacturer’s instructions. LNPs were diluted to a total lipid concentration of roughly 40 µg/mL prior to transfer into disposable polystyrene cuvettes (Sarstedt Inc., Newton, NC, USA) and read on the instrument.

#### De novo AML patient cohort analysis

Data from 1005 de novo AML specimens were obtained from 4 published cohorts: Beat AML (Bottomly *et al*., 2022), AML PMP (Docking *et al*., 2021), TCGA LAML (Cancer Genome Atlas Research *et al*., 2013), Leucegene.

(a) Preprocessing Gene expression count data were combined and batch corrected using Combat-seq, and filtered to remove genes with low expression (0 in ≥ 50% of patients) and low variance (standard deviation ≤ 0.5 across patients) (Zhang et al., 2020). Small variants (single nucleotide variants and indels) were called from RNA-seq data using GATK SplitNCigarReads followed by variant calling by GATK HaplotypeCaller, Vardict, Varscan, and Freebayes. Variants in the Canadian PanHeme Panel’s AML/MDS/MPN/CHIP/Various Heme Pathologies genelists (n = 99 genes) were retained if called by 2 or more callers (except for NPM1, which was retained if called by 1 or more caller), and were filtered to remove calls with < 7 supporting reads, strand biased calls, common population polymorphisms (gnomAD allele frequency > 0.01), known benign/likely benign ClinVar variants, synonymous variants without predicted splicing effects, or recurrent spurious calls. Fusion transcripts were called by STAR Fusion and Arriba and were filtered to fusions involving genes known to undergo recurrent fusions in AML.
(b) Gene expression analysis RNA-seq samples were divided into PAN3-AS1 high (top tertile) and low (bottom tertile) groups.

(i) Differential gene expression between groups was analyzed using the DESeq2 algorithm. Differentially expressed genes were compared between the 4 cohorts, to identify common significant up- and down-regulated genes.
(ii) Gene set enrichment analysis was performed between groups using GSEA software (Subramanian et al., 2005), using the Hallmarks and C2 Biocarta gene sets. Differentially expressed gene sets were compared between the 4 cohorts, to identify common significant differentially expressed gene sets.
(c) Expression vs. AML mutations

(i) Heatmap: Genes with sufficient sample size per group (mutated, unmutated) were included, to achieve 80% power to detect changes in gene expression between groups with effect size ≥ 0.8. Patients were arranged from lowest to highest expression of *PAN3-AS1*, and their mutation status per gene (mutated, unmutated) were plotted. Genes with significant differences in *PAN3-AS1* expression between mutated and unmutated patient groups are marked with asterisks (Mann-Whitney-U test with FDR correction, * p < 0.05, ** p < 0.01, *** p < 0.001).
(ii) Boxplots: For heatmap rows (genes) with a statistically significant difference in *PAN3-AS1* expression between mutated and unmutated patient groups, a boxplot was generated to visualize the *PAN3-AS1* gene expression between groups.

#### HiC data analysis

Hi-C data for MOLM13 cells (GSM4417606) were processed using Juicer tools, and topologically associating domains (TADs) were identified at 25 kb resolution with the Arrowhead algorithm. Differentially accessible ATAC-seq peaks were annotated as promoter, enhancer, or regulatory elements based on the ChromHMM segmentation for hematopoietic stem cells (HSCs) obtained from the Roadmap Epigenomics Project (https://egg2.wustl.edu/roadmap/web_portal/). TADs containing these differentially accessible peaks were extracted.

#### Quantification and statistical analysis

All the experiments were repeated with at least three biological replicates, unless otherwise specified in the figure legends. GraphPad Prism 10 software was used to generate graphs and to perform statistical analysis. P-values were calculated using student t-test, Kaplan–Meier method, or the appropriate statistical methods as reported in the figure legends. Results were shown as mean ± standard error of the mean (SEM). Statistical significance is determined by the value of p < 0.05. ns indicates not significant when the P-value is ≥ 0.05.

---

**Figure S1.**
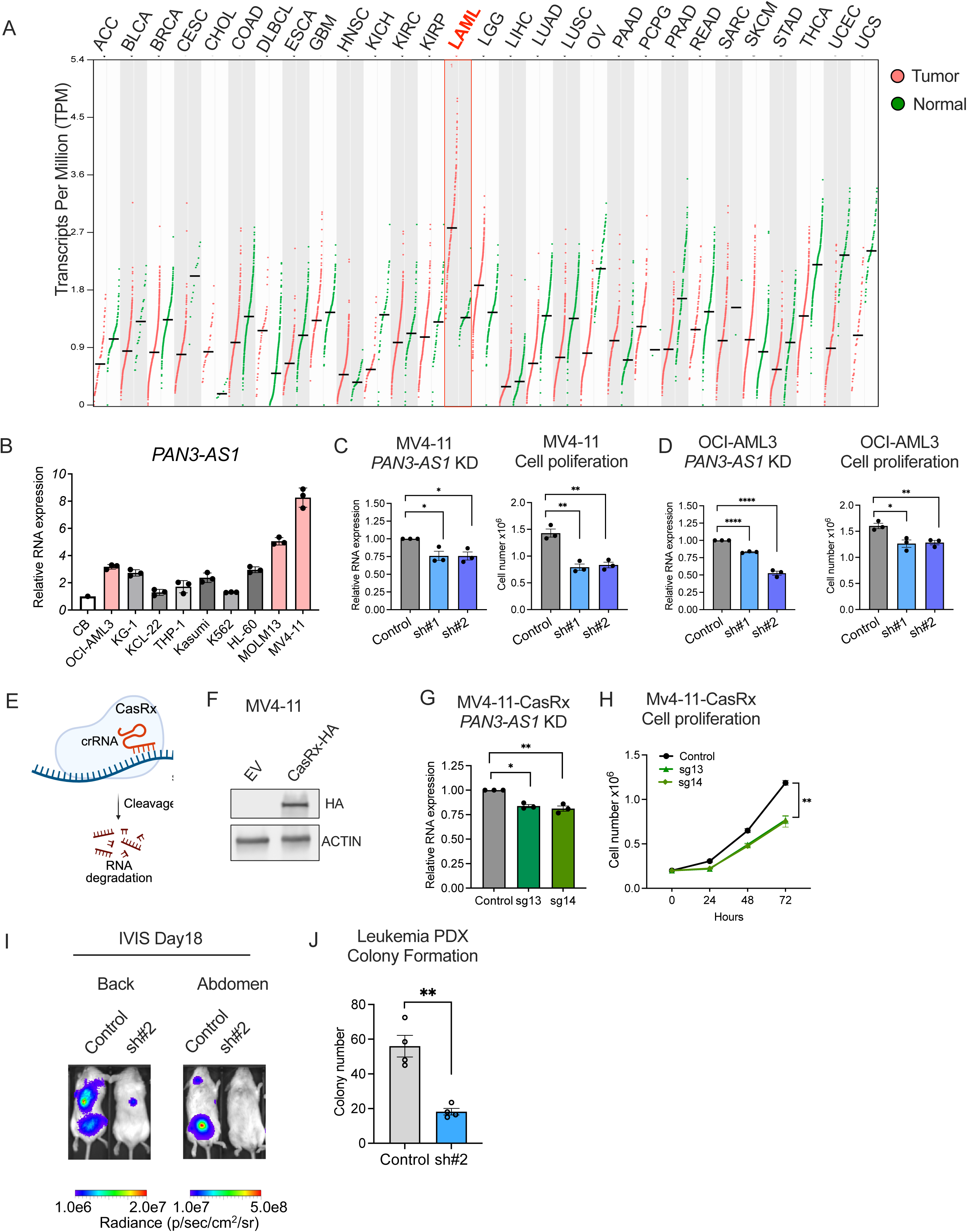

**Figure S2.**
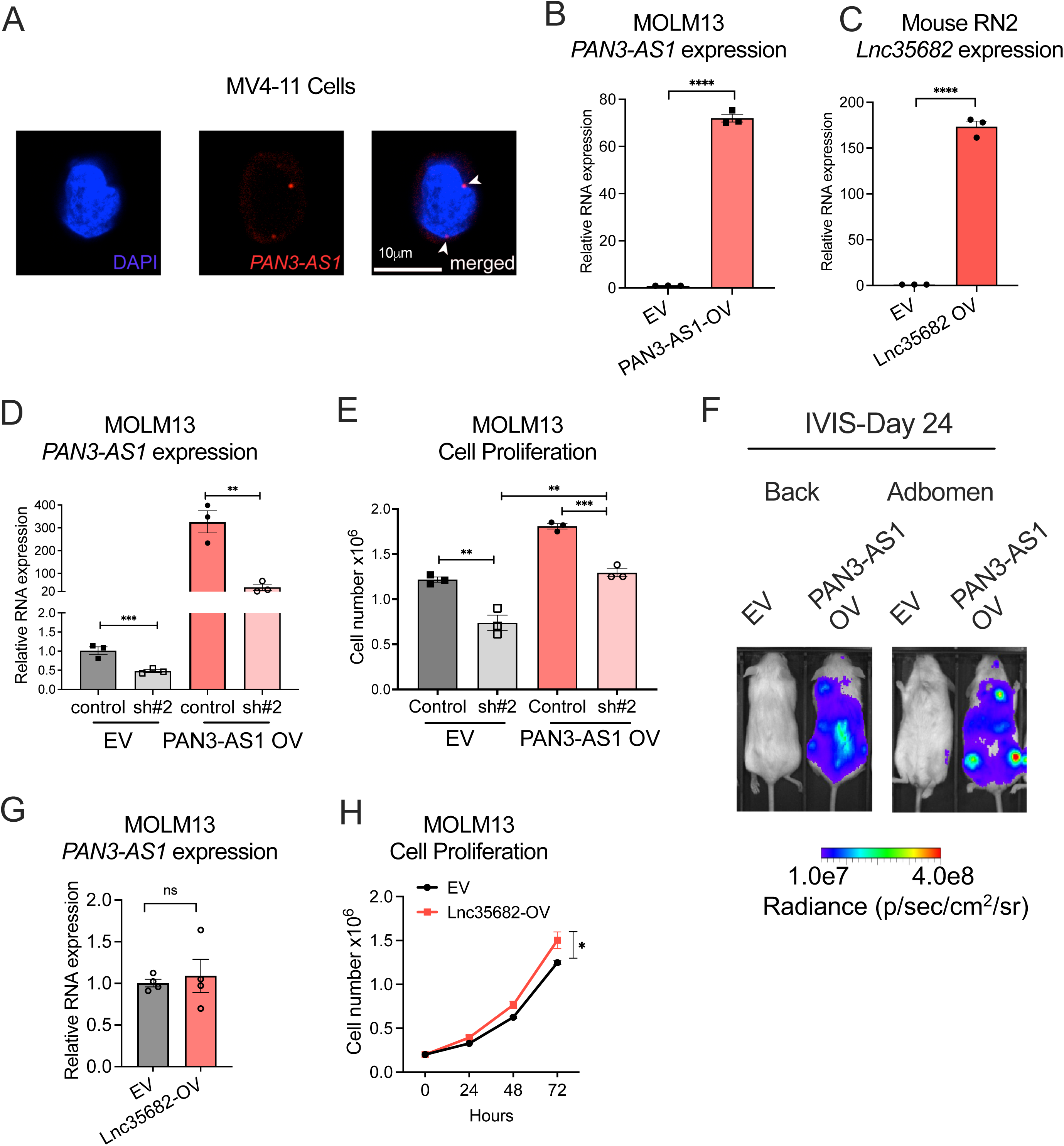

**Figure S3.**
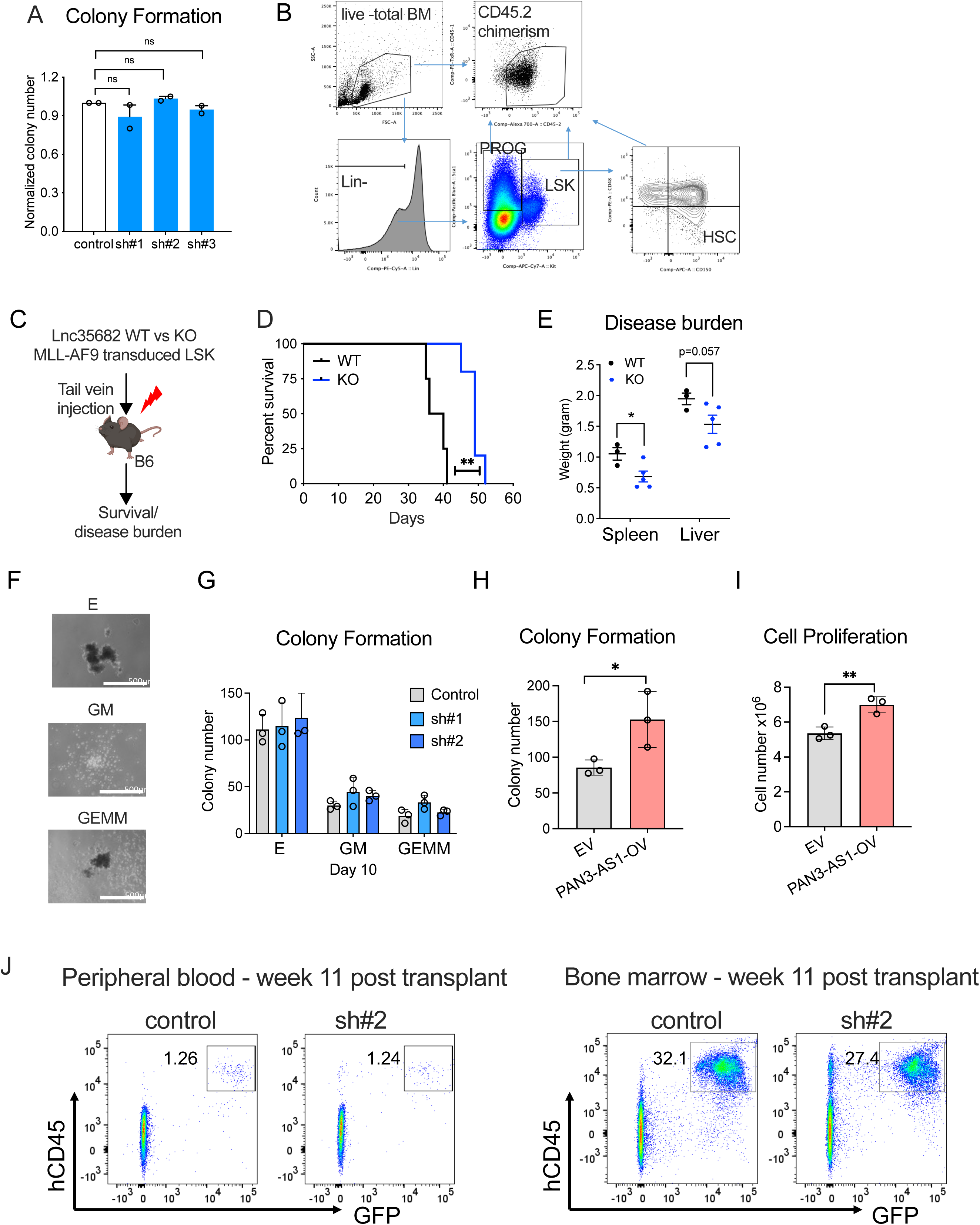

**Figure S4.**
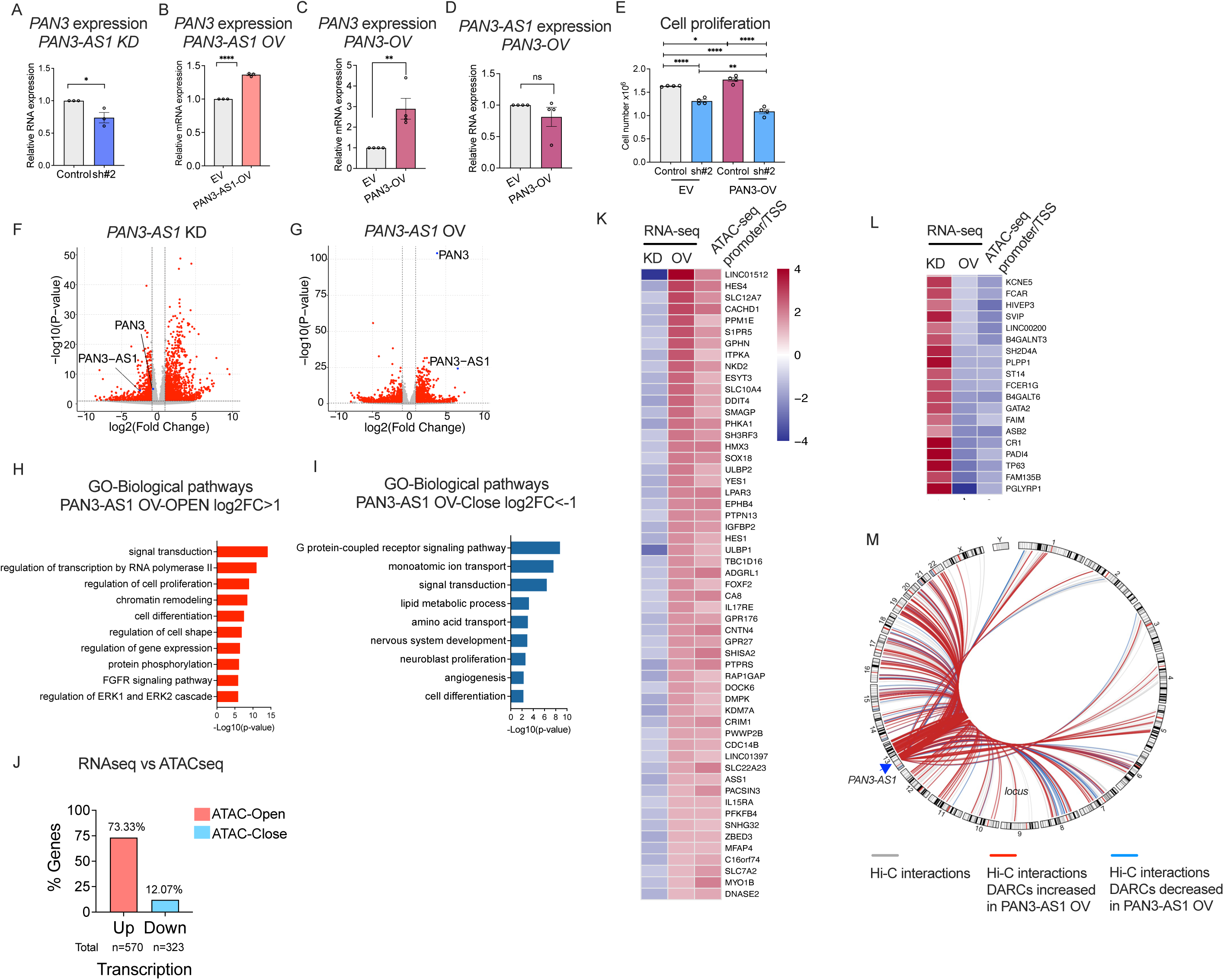

**Figure S5.**
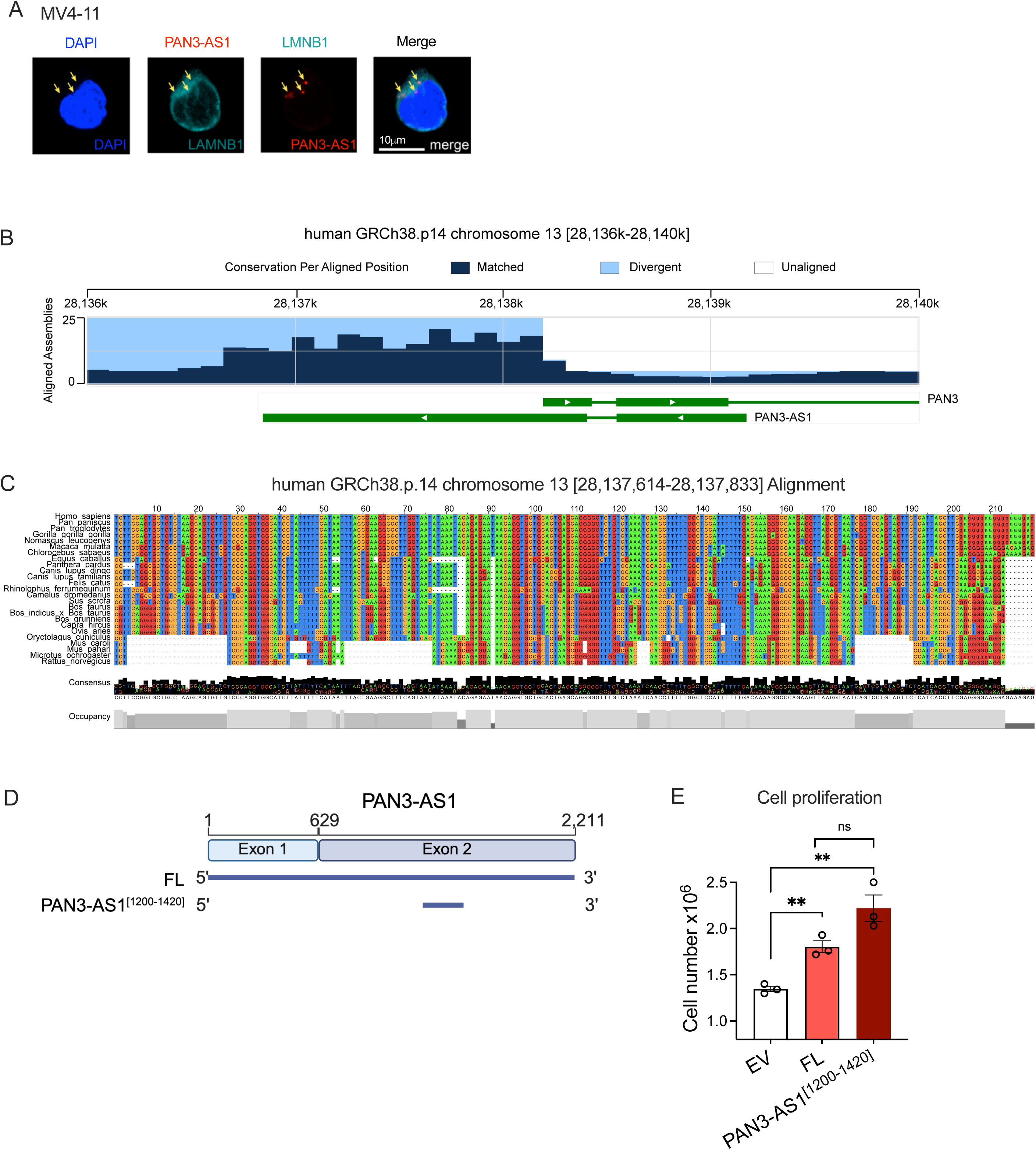

**Figure S6.**
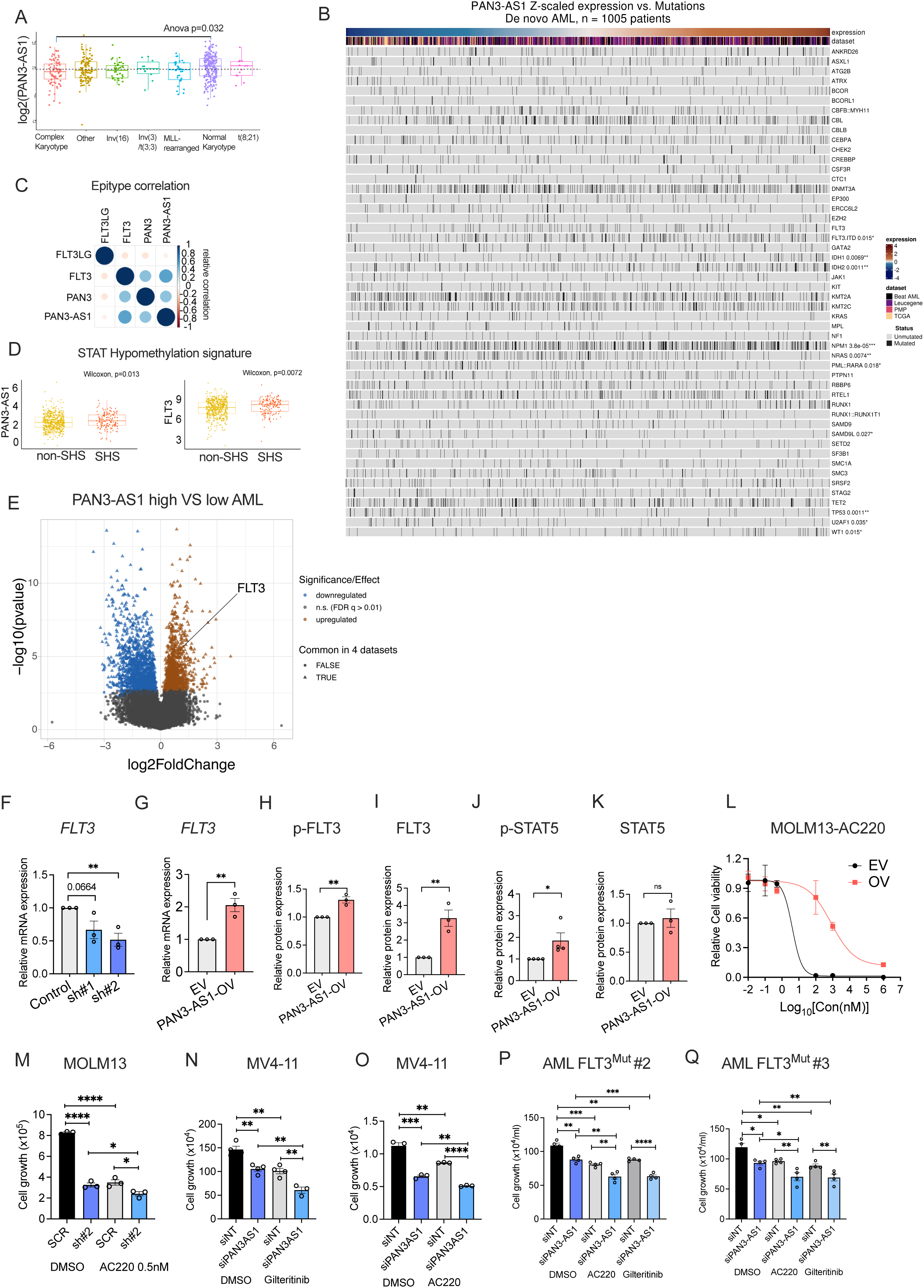

**Figure S7.**
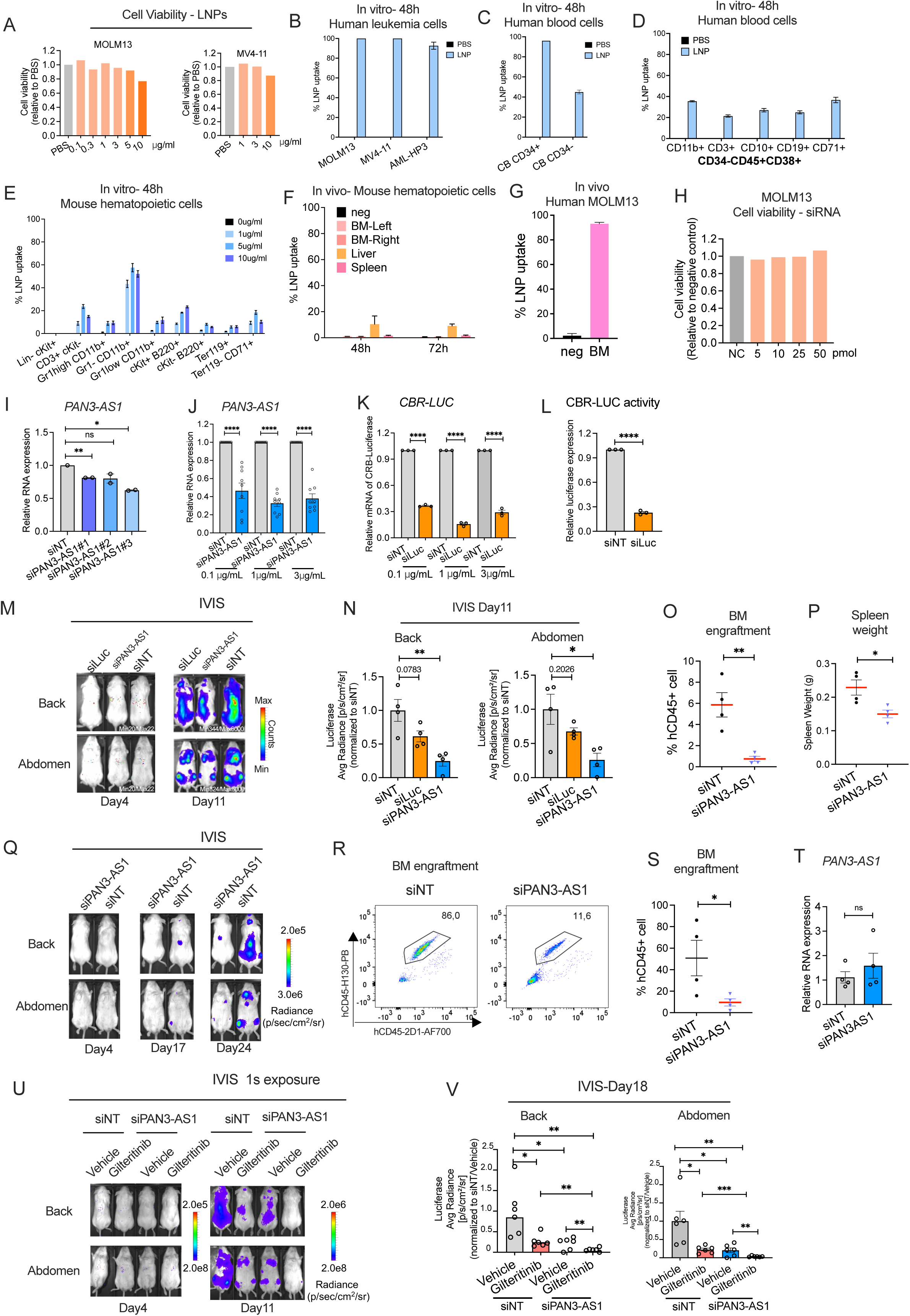

